# Flight performance of the highly invasive box tree moth *Cydalima perspectalis* (Lepidoptera: Crambidae)

**DOI:** 10.64898/2026.06.06.730098

**Authors:** Audrey Bras, Marie-Anne Auger-Rozenberg, Jérôme Rousselet, Alain Roques, Daniel Sauvard

**Affiliations:** URZF, INRAE – Orléans, France; Research Centre for Ecological Change, Organismal and Evolutionary Biology Research Programme, Faculty of Biological and Environmental Sciences, University of Helsinki, Helsinki, Finland

**Keywords:** *Cydalima perspectalis*, *Buxus*, Invasion, Insect, Flight mill, Flight capabilities

## Abstract

The multivoltine non-native box tree moth, *Cydalima perspectalis* (Lepidoptera: Crambidae), was accidentally introduced in several continents and exhibited a very rapid expansion in its invaded ranges, colonising around 40 countries in less than twenty years. Initially observed in urban areas, it quickly spread to forest stands, causing significant damage to boxwood trees. If the role of the ornamental plant trade in its fast invasion has been investigated, the insect’s flight capabilities has thus far been overlooked, limiting estimations of its natural dispersal. In this context, we assessed the flight performance of the adults and investigated potential effects of sex, age and generation using computer-monitored flight mills. Under these controled conditions, we estimated the distance flown per night and over the adults’ lifespan in both sexes for each generation. The adults were able to fly on average 18 km within their lifespan, even though distances flown were highly variable, including several females capable of performing long-distance flights (up to 150 km). The distances covered were partly correlated with the age and body mass of the adults. Mated females flew longer distances than virgin ones, but long dispersal seemed to limit their fecundity. Finally, the overwintering generation presented the highest flight capabilities with individuals able to cover 22 km on average while the late summer generation covered only 10 km. The box tree moth showed good dispersal abilities that likely played a significant role in its rapid local expansion, while human-mediated dispersal favoured its long-distance dispersal.

## Introduction

Biological invasions are recognised as a major component of global change, threatening biodiversity, public health and economy (Sakai et al., 2001; Simberloff et al., 2013). Global trade, travel and tourism have led to both intentional and accidental transport of many species outside their native range (Fenn-Moltu et al., 2022; Hulme, 2009; Perrings et al., 2005). In Europe, introduction of non-native species, especially insects, is still increasing without signs of saturation (Seebens et al., 2017). Furthermore, since about twenty years ago, a number of non-native insect species entering Europe have expanded very quickly after establishment, likely due to the liberalisation of trade and travel in the European Union, which may in turn have facilitated their propagation (Roques et al., 2016). Dispersal is a key factor in invasion processes (Blackburn et al., 2011; Renault et al., 2018; Wilson et al., 2009), affecting the degree of damage that a non-native insect can cause. Species self-mediated dispersal capabilities can facilitate establishment and expansion phases as individuals search for food, reproductive partners and new habitats (Bowler and Benton, 2005; Renault et al., 2025). However, human activities can provide vectors for dispersal which may likewise accelerate species spatial expansion after their introduction (Estoup and Guillemaud, 2010; Garnas et al., 2016). Thus, it can be challenging to distinguish between the contributions of non-native species own dispersal from human-mediated dispersal in the spread process (Essl et al., 2018). Hence, separating self-mediated dispersal from human-mediated dispersal is required to understand and predict both species expansion and distribution (Robinet et al., 2012, 2017) and to improve non-native species management (Robinet et al., 2019), especially in a context where species are spreading faster than before (Roques et al., 2016).

The box tree moth, *Cydalima perspectalis* (Walker, 1859) (Lepidoptera: Crambidae), is an Asian native Lepidopteran developing on boxwood trees from the genus *Buxus* (Kawazu et al., 2010a; Maruyama and Shinkaji, 1991; Wan et al., 2014). Naturally present in Japan, Korea and China, it was accidentally introduced in Europe in the early 2000s (Kenis et al., 2013), and in North America few years ago (Coyle et al., 2022). Its expansion in Europe is representative of the fast spread observed for non-native insects accidentally introduced after the 1990’s (Roques et al., 2016). Indeed, the moth was first recorded in 2007 simultaneously in Germany (Krüger, 2008) and in the Netherlands (van der Straten and Muus, 2010), from where it has quickly spread across Europe until Asia Minor, and is presently recorded from around 40 countries (Bras et al., 2025, 2019). *C. perspectalis* has from two to four generations per year, overwinters as young larvae in *Buxus* trees (Göttig and Herz, 2017; Nacambo et al., 2014) and presents two adult morph types: a dominant white morph and a melanic morph considered rare (Göttig and Herz, 2017). In several countries, the moth was first recorded in urban areas on ornamental boxwood trees and thereafter moved to natural forests, causing severe defoliation on native *Buxus* species such as *B. sempervirens, B. colchica* or *B. balearica* (Gil-Vives et al., 2026; Gninenko et al., 2014; Kenis et al., 2013; Mitchell et al., 2018). No efficient natural enemies are known in Europe, leading to population outbreaks and *Buxus* mortality as a result from repeated attacks by larvae feeding on leaves and bark of trees (Casteels et al., 2011; Nacambo et al., 2014). Thus, this insect represents a threat for *Buxus* trees commonly planted as ornamentals in historical, public and private gardens as well as for natural stands in southern Europe and in the Caucasus region (Di Domenico et al., 2012; Kenis et al., 2013; Mitchell et al., 2018).

The box tree moth was probably introduced in Europe with infested *Buxus* shipments (EPPO, 2012; Kenis et al., 2013), likely coming from Eastern China (Bras et al., 2022). Following at least three independent intercontinental introductions, the insect was accidentally transported across the continent through the intra-continental ornamental plant trade (Bras et al., 2019, 2022; EPPO, 2012), which could account for its fast spread. Several surveys on its local expansion in Europe (Bakay and Kollár, 2018; Kazilas et al., 2021; van der Straten and Muus, 2010) have however suggested that the moth’s active dispersal might have also played a significant role in its establishment and expansion. A population dynamics model showed that the local coexistence between the box tree moth and its host plant were only leading to extinction when long-distance dispersal was combined with a slow-growth of the host plant (Ledru et al., 2022). Few studies have estimated the natural dispersal of the moth to be about (5 to 10) km per year (Casteels et al., 2011; van der Straten and Muus, 2010) or (1 to 2) km per generation (Schmera and Baur, 2024). Nevertheless, if the flight activities of this nocturnal species have been assessed, showing a peak of activity two hours after the beginning of darkness and a decrease two hours before sunset (Kawazu et al., 2010b), the flight performance of the box tree moth has not yet been measured.

Characterisation and comparison of flight behaviour of wild insects is challenging in the field. Hence, several techniques have been developed to study the potentialities of insect dispersal, such as computerised flight mills. They allow the estimation of flight performance of a tethered insect under controlled conditions in the laboratory (Bruzzone et al., 2009; Minter et al., 2018). Species dispersal is a complex process influenced by individual conditions, behaviour, and several environmental stimuli (Bonte et al., 2024; Cote et al., 2017). Results from flight mill experiments thus need to be considered with caution, as they only measure individual physical abilities to flight (Le Souchu et al., 2025; Sauvard et al., 2018). This method has nevertheless several advantages, notably its ability to compare flight performance under different conditions or to assess the influence of different traits on flight capabilities such as wing color (Davis et al., 2012), age (Coombs, 1997), sex (Hughes and Dorn, 2002) or mating status of adults (Schumacher et al., 1997). This technic has been widely used to assess the flight performance of other Lepidoptera species (e.g., Hughes and Dorn, 2002; Qin et al., 2018; Shirai, 2006; Yang et al., 2017), or non-native species (e.g., Hoddle et al., 2015; Lopez et al., 2017; Sarvary et al., 2008a; Sauvard et al., 2018). Thus, despite shortcomings, flight mills are useful tools for defining the maximum flight capabilities of a given species and to acquire precise data on its flight activity.

In this study, we assessed the flight performance of the non-native box tree moth to better understand the insect’s role in its expansion. For this purpose, we tested on a computerised flight mill each generation of several close by invasive French populations of *C. perspectalis*, and this for both sexes. We characterised the moth’s flight capabilities and investigated potential effects of sex, age and generation. As reproduction can affect population establishment, we tested for potential trade-offs between flight capabilities and fecundity. Considering its fast expansion (Bras et al., 2019), we hypothesise that the flight capabilities of the moth are greater than previously reported (Casteels et al., 2011; Schmera and Baur, 2024; van der Straten and Muus, 2010) while steady across generations. We predict that female would have greater flight abilities than male, and that mating will have a positive effect on their flight. The results will improve our understanding on the role of the box tree moth’s active dispersal in its invasion in Europe and participate in developing more efficient strategies to control this pest threatening *Buxus* species (Coyle et al., 2022; Mitchell et al., 2018).

## Material and methods

### Insect collection and rearing

All insect collections were carried out in urban areas during the years 2017 and 2018 in the French region Centre-Val de Loire, where the first detection occurred in 2014. In 2017, collections were carried out in Saint-Denis-sur-Loire and incidentally in Mettray for the over-wintering generation (G1), in both Olivet and Lamotte-Beuvron for the second generation (G2), and in Lamotte-Beuvron for the last generation (G3) (Table 1). In spring 2018, supplement collections were carried out in Azay-le-Ferron, Bouges-le-Château, Bucy-Saint-Liphard and Montlouis-sur-Loire (overwintering generation, E2018). For each generation, last instar larvae were hand-collected on ornamental boxwood trees. Sampled insects were brought back to the laboratory, put in 29 cm *×* 28 cm *×* 9 cm plastic boxes with boxwood tree branches, and reared until the end of pupation at (20 *±* 1) ^*°*^C, with a photoperiod of 16:8 LD and 70 % of humidity. Adult emergences were checked every morning, and emerged moths were taken out of the box. Adults presenting unexpanded or wrinkled wings were not tested. White morph adults were chosen randomly and prepared for experimentations. Adults showing the rare melanic morph were not used, except as mating partners, if needed. Males and females were separated in order to avoid mating, labeled and kept in 30 cm *×* 30 cm *×* 30 cm mesh cages under the same rearing conditions as larvae and pupae, with unlimited food (cotton saturated with a 30 % honey solution) and water.

**Table 1.**
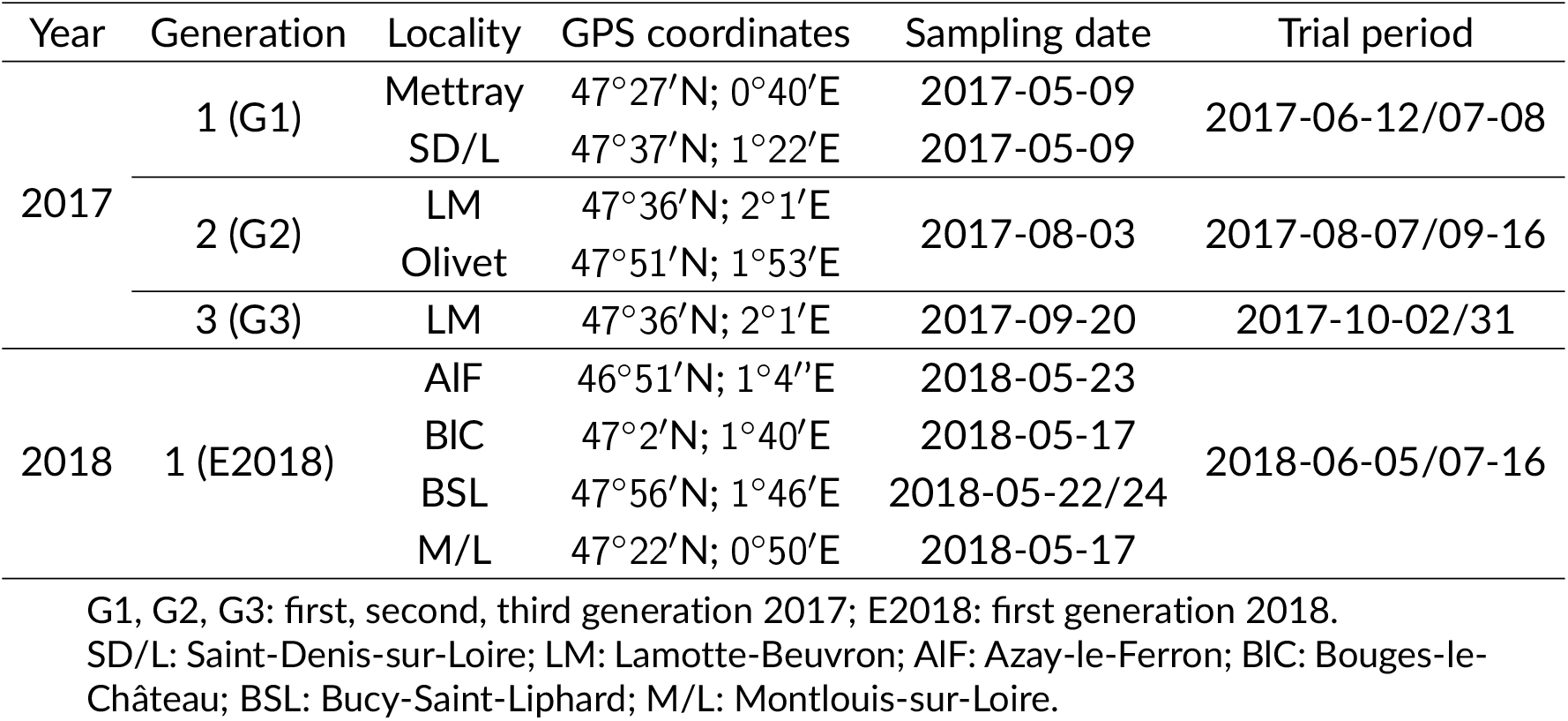
Description of *Cydalima perspectalis* samples used in flight mill experiments. All localities are in France, in Centre-Val de Loire region.

### Flight mills

The flight mill setup matched the design described by Sauvard et al. (2018). Briefly, the base of the flight mill was a cast iron support stand into which was screwed a threaded shaft. A hole was drilled into the top of the shaft, into which a 3-mm-diameter rod was inserted. A carbon-fibre arm was then attached to a miniature ball bearing, which was placed around the rod. This setup minimized friction and allowed the arm to rotate easily. A small foam block was glued to one end of the arm. Two tiny pins were inserted into the foam. They were used to secure the insect tether to the apparatus, so insect could drag the arm when it flew. The foam was positioned along the arm such that an insect travelled 2 m per lap. The movement of the carbon-fibre arm was monitored using a computerised system based on an infrared emitter and an associated detector, both connected to a PCI board installed in a computer. Triangular pieces of paper were taped to the centre of the arm to trigger the motion detection system. When the arm cut the beam between the emitter and the detector, an electrical signal was sent to the PCI board. An application written in C++ detected and time-stamped these signals and wrote the resulting data into a log file. Insect flight activity was thus precisely recorded. The flight mills were kept in the laboratory and placed on shelves. They were located in three separated rooms. A computer with its PCI board could monitor up to eight flight mills. Thirty-two computerised flight mills were available to test insect flight capabilities for G1 and G2, and 16 for G3 and E2018. As the box tree moth is nocturnal, lights were switched off during flight tests.

The insects were tethered to the flight mill using a tether made with a piece of Bristol paper at one end of which a small piece of foam was glued with cyanoacrylate glue (Diall®). After emergence, insects were labeled and put at 3 ^*°*^C for one hour. After this time, insects were inactive and scales on the thorax were removed. The tether was then glued on top of the thorax using neoprene adhesive gel (Diall®), with the foam part placed above the insect’s head. Precautions were taken to avoid putting glue on insect wings. The moths were tethered to the flight mill by plugging tether foam on the arm pins. The tether was left on the thorax until the insect’s death. If the tether was loose, it was glued again on the insect’s thorax following the same procedure as explained above.

The flight tests were performed at room temperature (range: (20 to 25) ^*°*^C). Each tested insect was beforehand weighed, then placed on the flight mill while recording simultaneously started. During the flight test, no attempt was made to stimulate flight, and no food was provided. At the end of the flight test, recording was stopped, the insect removed from the flight mill and weighed again to estimate its mass loss. It was returned to the rearing room and immediately fed with water and honey solution. To minimize a possible room effect, insects were transferred to another flight mill room every other test. A small number of flight tests experienced technical problems, mainly insect detachment from the arm or power supply failure, leading to incomplete or missing records. Thus, these tests were removed from further analyses, except for values aggregating the results of several tests, which could then be slightly underestimated.

### Experimental design

We first aimed to characterise *C. perspectalis* flight capabilities. To do so, two modalities of weekly test design were used for individuals collected in 2017. The first modality, called A-Normal thereafter, was as follows: two consecutive test nights, one night of rest, two consecutive test nights, and two consecutive nights of rest. The second modality, called B-Intensive, was as follows: five consecutive nights of tests followed by two nights of rest. During the rest periods, each adult was left in a mesh cage with water and sugar under rearing conditions. As we were interested in assessing the effect of reproduction on the box tree moth’s flight capabilities, a subset of the females from the A-Normal experiment was allowed to mate. The selected females were at least 3 days old, so they had completed two test nights. Prior being allowed to mate, they were marked on the forewing in case they lose their tether. Afterwards, females were left in 30 cm*×*30 cm*×*30 cm mesh cages with untested males during two consecutive nights and the day in between with food and water. As competition seems to favour mating (authors personal observation), three to four females were placed in the cage with double the number of males. After the mating period, each female was kept in separate cages when not on the flight mill, and the presence of eggs in each cage was checked daily. This subset of females was called “mated”, while the rest was called “virgin”. Whatever the modality, insects of both sexes were tested from the night after their emergence until their death. The flight tests were done during the night from the end of the day until the early morning, and lasted about (16 *±* 2) hours. As *C. perspectalis* natural dispersal is directly dependent on the female flight capabilities, females were given priority for testing. Due to material constraints and variability in adult emergences during the experiments, the weekly test designs were slightly adjusted when needed (mostly one-day shifts, or, for mating, a moving after the third or the fourth test nights). Table 2 indicates the number of involved insects and carried out tests.

**Table 2.** Number of individuals (in **bold**) and tests (in roman) for both sexes of *Cydalima perspectalis* that were tested on a flight mill for each generation in 2017 experimental design. Numbers of tests used for further analyses are indicated in parentheses (Others were excluded due to technical problems).A, B: A-Normal, B-Intensive modality; G1, G2, G3: first, second, third generation; F, M: females, males.

|  |  | G1 |  | G2 |  | G3 |  | Total |
| --- | --- | --- | --- | --- | --- | --- | --- | --- |
|  |  | F | M | F | M | F | M |  |
| A | Mated | **8** | **0** | **17** | **0** | **4** | **0** | **29** |
|  |  | 52 (49) | 0 (0) | 117 (116) | 0 (0) | 22 (22) | 0 (0) | 191 (187) |
|  | Virgin | **17** | **9** | **30** | **15** | **14** | **9** | **94** |
|  |  | 52 (50) | 25 (24) | 124 (120) | 56 (56) | 50 (49) | 24 (24) | 331 (323) |
| B |  | **16** | **6** | **30** | **15** | **11** | **5** | **83** |
|  |  | 71 (68) | 15 (15) | 152 (149) | 50 (49) | 48 (48) | 21 (21) | 357 (350) |
| Total |  | **41** | **15** | **77** | **30** | **29** | **14** | **206** |
|  |  | 175 (167) | 40 (39) | 393 (385) | 106 (105) | 120 (119) | 45 (45) | 879 (860) |

We secondly aimed to analyse the effect of stress induced by flight tests on fecundity and determine potential trade-offs between flight duration and reproduction. For this purpose, only females were tested in 2018. They were randomly assigned to one of the six different weekly flight test profiles: 4 night tests a week (as in 2017 A-Normal modality), 2 night tests a week, 1 night test a week, 2 short tests a week, 1 short test a week, or control in which females were never tested. Night tests were carried out as in 2017 experiment. Short test designs were similar, but shorter timewise; they began at the end of the day and were stopped during the night, so that they lasted about 4 hours. Consequently, the weekly profiles represented a stress of either 64, 32, 16, 8, 4 or 0 hour(s) of flight tests per week, respectively. All females were allowed to mate, following the same protocol as in 2017. After mating, they were returned to their corresponding flight test design, and kept in separate cages between the flight tests. Presence of eggs was checked daily and they were counted if any. Beside control females simply maintained during their entire lifetime in the rearing room, referred thereafter as Base controls, three additional non-flying control protocols were setup. The first one was to test the effect of host plant stimulation on female’s egg laying. For this, a boxwood tree branch was added in the cage of some females maintained in the rearing room (Boxwood tree controls). The second control protocol was designed to check for any effect of the tether on insect (e.g., through its mass). Thus, a tether was glued on the thorax of some females which were kept in the rearing room until they died (Tether controls). Finally, the last protocol was used to test potential negative effects of fasting on female’s lifespan and fecundity. A tether was glued on the thorax of some females; they were maintained in the rearing room, except 4 nights a week to simulate flight test conditions of A-Normal experiment. During these nights, they were isolated in test rooms without food and water for the duration of a flight test (Fasting controls). Table 3 indicates the number of involved insects and carried out tests.

**Table 3.** Insects involved in 2018 experimental design: number of females of *Cydalima perspectalis* (in **bold**) that were either tested on a flight mill, with corresponding number of tests (in roman), or attributed to one of the four control modalities. Stress was expressed as flight test hours per week. Numbers of tests used in statistics are indicated in parentheses (Others were excluded due to technical problems). Control modalities: Base) insect left in rearing room; Boxwood) *idem* Base with a boxwood branch added; Tether) *idem* Base with tether glued on insect; Fasting) insects with tether in test room during 4 nights a week, without food or water. Only the first generation in 2018 was tested.

|  | Stress (h/w) | 4 | 8 | 16 | 32 | 64 | Total |
| --- | --- | --- | --- | --- | --- | --- | --- |
| Flight test | Number of females | **9** | **9** | **25** | **12** | **11** | **66** |
|  | Number of tests | 13 (13) | 34 (34) | 36 (36) | 34 (34) | 39 (37) | 156 (154) |
| Control | Modality | Base | Boxwood | Tether | Fasting |  | Total |
|  | Number of females | **16** | **14** | **4** | **4** |  | **38** |
| Total number of females |  |  |  |  |  |  | **104** |

### Data analysis

Ruby scripts were developed to analyse the logs recorded by the flight mill system. The log from each flight test was cleaned and divided into different phases. Flight phases occurred when moths were continuously moving, and they were separated by rest phases, defined as no movement for at least 5 s. For each test, the scripts determined the number and type of phases. For each phase, they computed duration, and, for flight ones, flown distance and mean speed. Ruby and R scripts then summarized these data for each test ((test) total flight duration and distance, mean and maximum flight phase duration and distance, and mean and maximum flight speed) and each insect ((lifetime) flight duration and distance). We also used insect data associated with each test (age, pre-flight and post-flight body mass, and so computed (absolute) mass loss and relative mass loss) or each insect (lifespan and fecundity).

All statistical analyses and figures were performed using R software (Version 4.2.2; R Core Team, 2022), with 95 % confidence level. As most data distributions were non-normal (often highly skewed without simple transformation to normalise them), we used nonparametric methods: Kruskal-Wallis rank sum test (followed by Dunn test with the Holm adjustment method when there were more than two groups) to compare group positions, and Spearman rank correlation for variable correlations. Difference between proportions were tested using Pearson’s chi-squared test. As each moth was tested several times, statistical tests could be biased due to autocorrelation. To deal with this issue, tests were mostly done on only the first flight test of the insects, or on values summarised by insect.

Abbreviations used in text: KW: Kruskal-Wallis rank sum test; Dunn: Dunn’s test for pairwise multiple comparisons of the ranked data; Spearman: Spearman rank correlation (*ρ*: Spearman rank correlation coefficient); *χ*^2^: Pearson’s chi-squared test; *M*: arithmetic mean; *SD*: standard deviation; *Mdn*: median; *Q3*: third quartile; *Max*: maximum; *p*: *p*-value; *R*^2^: adjusted R-squared; NS: not significant.

## Results

We compared the different flight performance values between sampling locations for each generation, but no significant difference was found; we thus considered males and females without distinguishing between sites. Similarly, no significant difference in flight capabilities was found in 2017 between the two experimental designs A-Normal and B-Intensive, so flight characteristics are thereafter provided without experimental distinction. In flight capability experiment (2017), we tested 147 females and 59 males on computerised flight mills (Total 206; Table 2). In trade-off experiment (2018), we tested 66 females on computerised flight mills, leaving out 38 as controls (Total 104; Table 3).

### Body mass and flight capability during flight tests

In flight capability experiment (2017), 879 flight tests were carried out in total (G1: 215; G2: 499; G3: 165; Table 2). Excluding mated females (see below), females were slightly more tested on flight mill than males (*Mdn* = 3 for both sexes, but *M* = 4.2, *Q3* = 5 and *Max* = 16 with females, while *M* = 3.2, *Q3* = 4 and *Max* = 13 with males; KW *p* = 0.003). No significant difference was found between treatment nor generation.

Insect pre-flight body mass was variable, ranging from 30 mg to 232 mg (90 % between 54 mg and 130 mg) with a mean value of 89 mg for females and 74 mg for males. Females were slightly heavier than males. In the first test, this was only significant for G2 (KW *p* < 0.001). Body mass of both sexes tended to decrease from G1 to G2 and especially G3. In the first test, this was significant for females (KW *p* < 0.001; males KW *p* = 0.091; Figure 1, Table S1 in Sauvard et al. (2026b)). Insect body mass slightly decreased with insect age, (Females: Spearman *p* < 0.001, *ρ* = *−*0.14; males: Spearman *p* < 0.001, *ρ* = *−*0.54. Figure 2).

**Figure 1.**
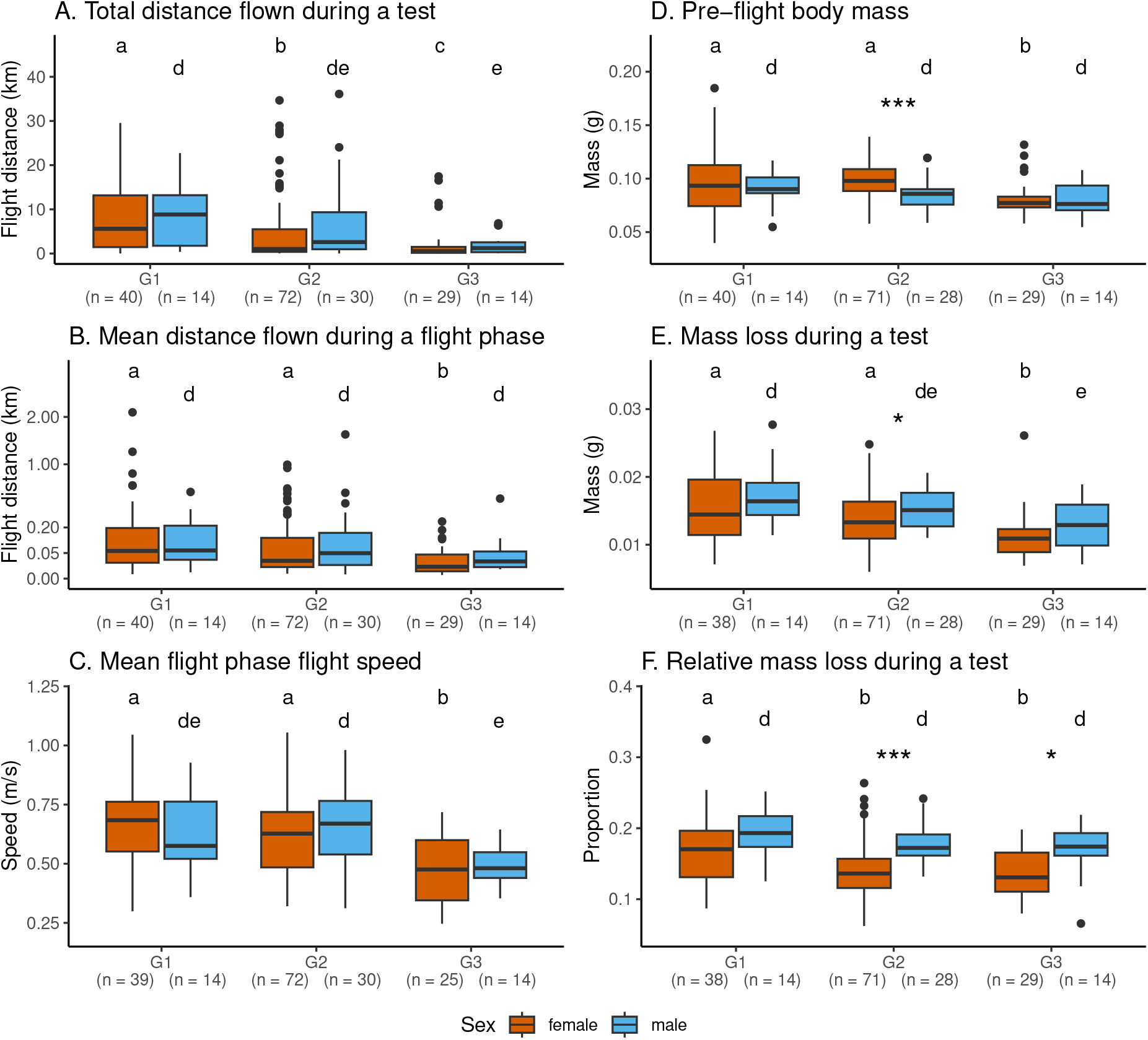
Flight parameters for *Cydalima perspectalis* for each generation and sex. Experiment 2017, only first test of each insect. The modalities A-Normal and B-Intensive were mixed as no difference was observed. Mean distance flown during a flight phase is displayed with square root scale to improve visibility. Significant differences between generations: 1) global KW A female *p* < 0.001, A male *p* = 0.024, B female *p* = 0.002, B male NS, C female *p* < 0.001, C male *p* = 0.007, D female *p* < 0.001, D male NS, E female *p* < 0.001, E male *p* = 0.034, F female *p* = 0.005, F male NS; 2) pairwise differences are indicated by letters (Female: abc, male: de; Dunn *p* < 0.05). Stars, if any, indicate a significant difference between sexes (KW, *: *p* < 0.05, ***: *p* < 0.001).

**Figure 2.**
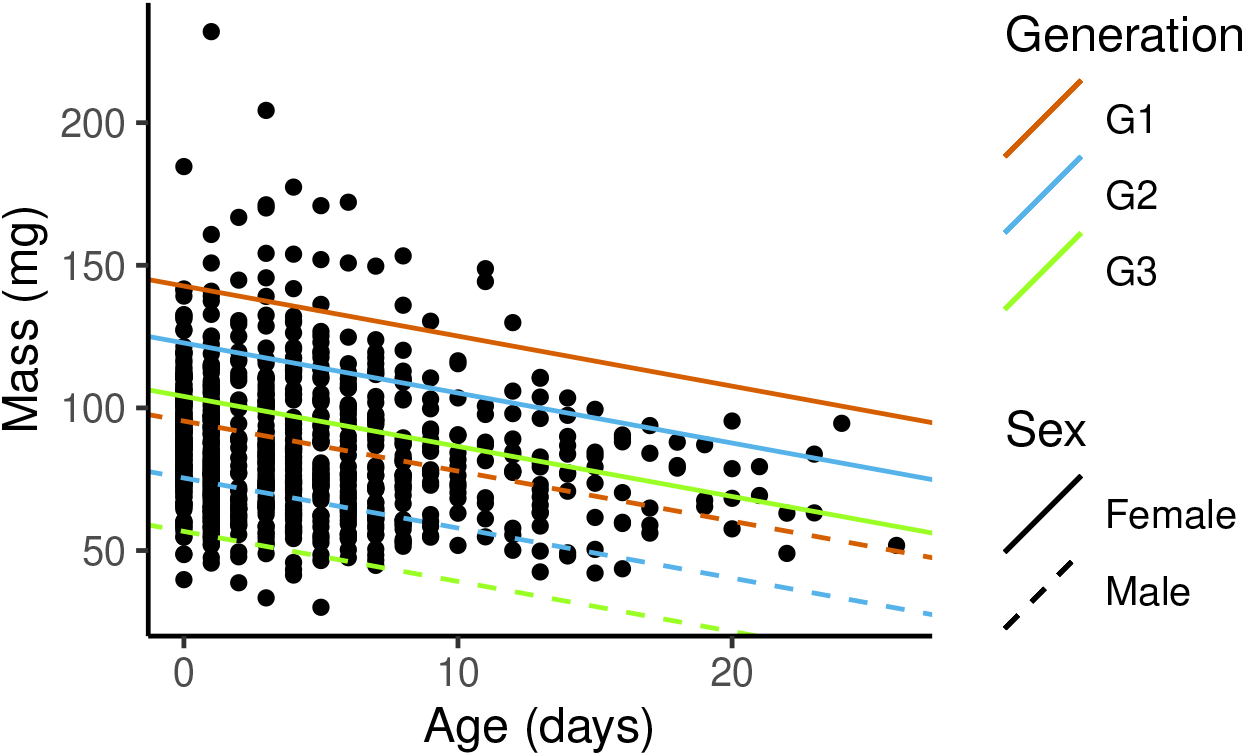
*Cydalima perspectalis* mass at the beginning of a flight test according to the insect age. Experiment 2017 (all tests gathered). Global correlations (Spearman): females *p* < 0.001 *ρ* = *−*0.14, males *p* < 0.001 *ρ* = *−*0.54. Group lines were extracted from an approximate linear model analysis: *mass ∼ age + sex + generation + insect_id* (*p* < 0.001, *R*^2^ = 0.76), where a large part of the variability is due to individual effects (*insect_id*), especially for females.

Distance flown during a test was highly variable ((0 to 45) km). Its distribution was also extremely skewed, as most insects flew a short distance, while a few covered far longer distances (mean 4.5 km, but median only 1.1 km; Figure 3). Distribution shape was similar in all subgroups (e.g., by experimental designs or generations). Even if flight distance medians for males were longer than those for females in the first test, these differences were not significant (Figure 1, Table S1 in Sauvard et al. (2026b)). On the other hand, distance flown during a test decreased from G1 to G3 in both sexes (KW in the first test: females *p* < 0.001, males *p* = 0.024). The evolution of flight capabilities with insect age depended on insect sex and mating status (Figure 4). For both sexes, only youngest insects (especially those five days old or less) were flying the longest distances. However, average flown distances declined less sharply than these maximums, especially for mated females (Spearman for virgin or mated females: NS; for males: *p* = 0.02, *ρ* = *−*0.17. Figure 4). Oldest females thus tended to fly longer than oldest males, but this difference remains unsignificant. During a flight test, insects that flew performed a highly variable number of flight phases (1 to 956, *Mdn* = 49). Distance flown during a flight phase was also highly variable and, in most cases, very short (Per-test mean distance: (0.001 to 6.83) km, *Mdn* = 0.017 km; per-test maximum distance: (0.001 to 26.3) km, *Mdn* = 0.275 km). Per-test mean and maximum flight phase distances were correlated with total distance flown during a test (Spearman *p* < 0.001, *ρ* = 0.88, and *p* < 0.001, *ρ* = 0.94, respectively). Consequently, they varied rather similarly with sex and generation (Figure 1, Table S1 in Sauvard et al. (2026b)).

**Figure 3.**
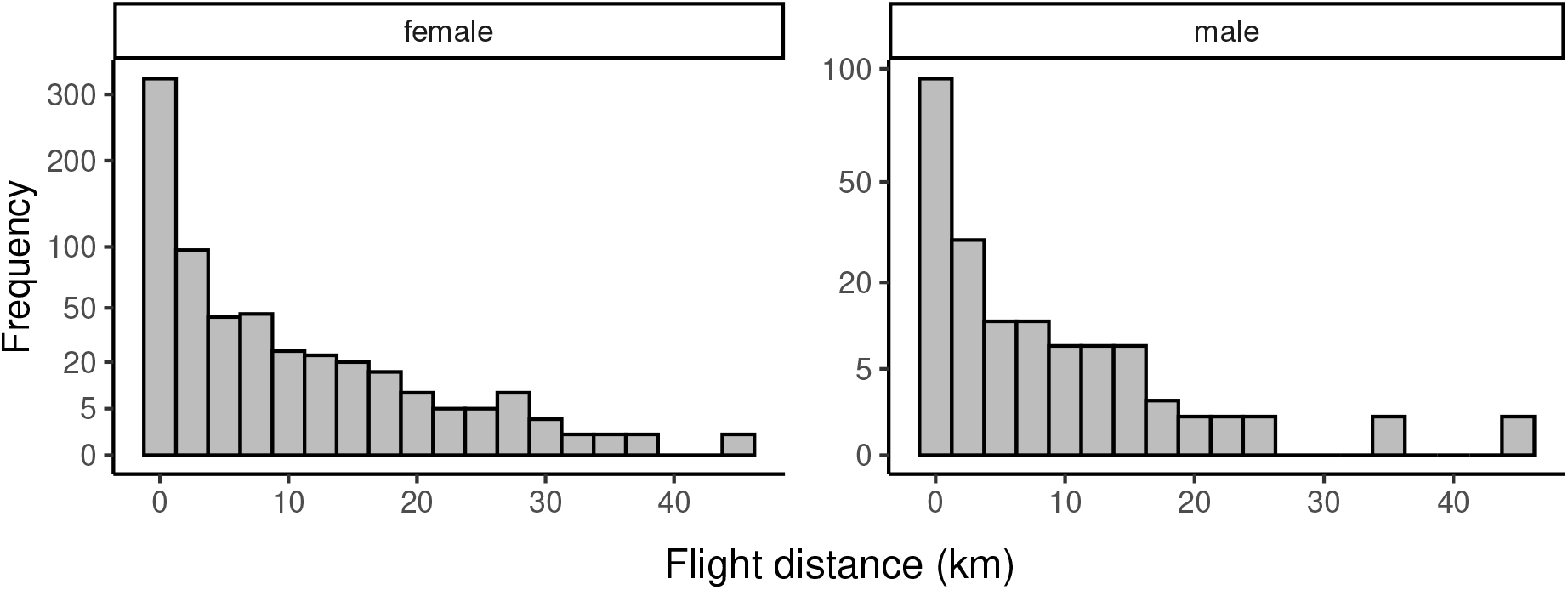
Distribution of distances flown during a flight test for both sexes of *Cydalima perspectalis*. Experiment 2017. No distinction between modalities and generations was done as distributions were showing similar shapes. Frequency is displayed with square root scale to improve visibility.

**Figure 4.**
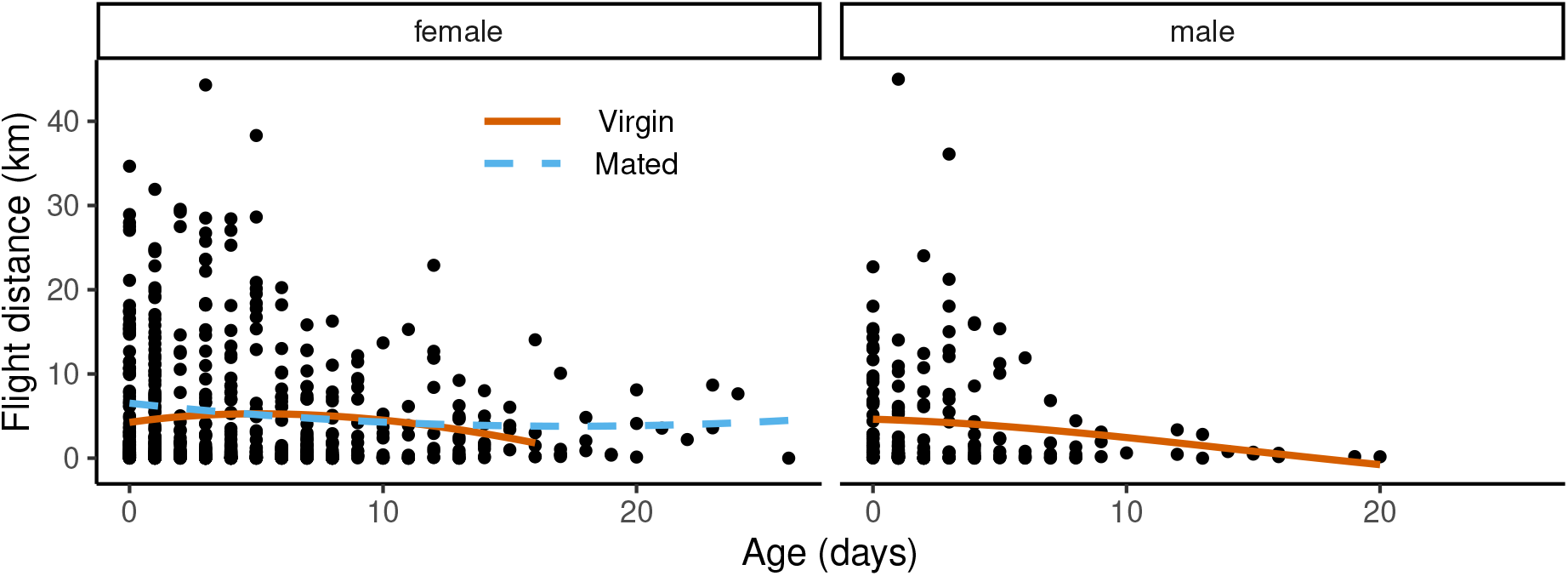
Distances flown per flight test according to age and sex for *Cydalima perspectalis*. Experiment 2017 (all tests gathered). Global correlations (Spearman): virgin or mated females NS, males *p* = 0.02 *ρ* = *−*0.17. Colored lines are loess smoothings of dot clouds (Virgin and mated females are separated).

Distance flown during a test was loosely related to the insect’s pre-flight body mass (Figure S1 in Sauvard et al. (2026b)). For females, the longest distances were covered by those of median body mass. The pattern was less consistent for males. In G1, distance flown was positively correlated with insect’s pre-flight body mass (Spearman *p* < 0.001, *ρ* = 0.63), while in G2 and especially G3, this pattern in males became similar to the one found in females.

Insects’ flight induced body mass loss (Figure S2 in Sauvard et al. (2026b)). Body mass loss and relative body mass loss were correlated with flown distances (Spearman *p* < 0.001, *ρ* = *−*0.48, and *p* < 0.001, *ρ* = *−*0.38, respectively). Globally, body mass loss, and especially relative body mass loss, were slightly greater for males than for females, but the differences were not always significant (Figure 1, Table S1 in Sauvard et al. (2026b)). Body mass loss also decreased from G1 to G3, especially for females; this decrease was less pronounced with relative body mass loss, and was not significant for males (Figure 1, Table S1 in Sauvard et al. (2026b)). Insect age did not affect body mass losses.

Insect speed on flight mills was much less variable than flight distance. It was not affected by sex, but it decreased from G1 to G3 (Figure 1, Table S1 in Sauvard et al. (2026b)). Independantly, mean flight speed was positively correlated with total flight distance (Spearman *p* = 0.007, *ρ* = 0.37, *p* < 0.001, *ρ* = 0.48, and *p* = 0.002, *ρ* = 0.48, in G1, G2 and G3, respectively). Mean flight speed also decreased with age, except in G3 (Spearman *p* = 0.005, *ρ* = *−*0.2, *p* < 0.001, *ρ* = *−*0.36, and *p* = 0.433, *ρ* = *−*0.07, for G1, G2 and G3, respectively).

### Adult lifespan and flight capability

Tested insect lifespan was highly variable (1 to 27 days, *Mdn* = 5 days). It did not vary between generations, but it depended on sex and mating status (KW *p* < 0.001): mated females survived longer than virgin ones while the latter had a slightly longer lifespan than males (all virgin) (Figure 5).

**Figure 5.**
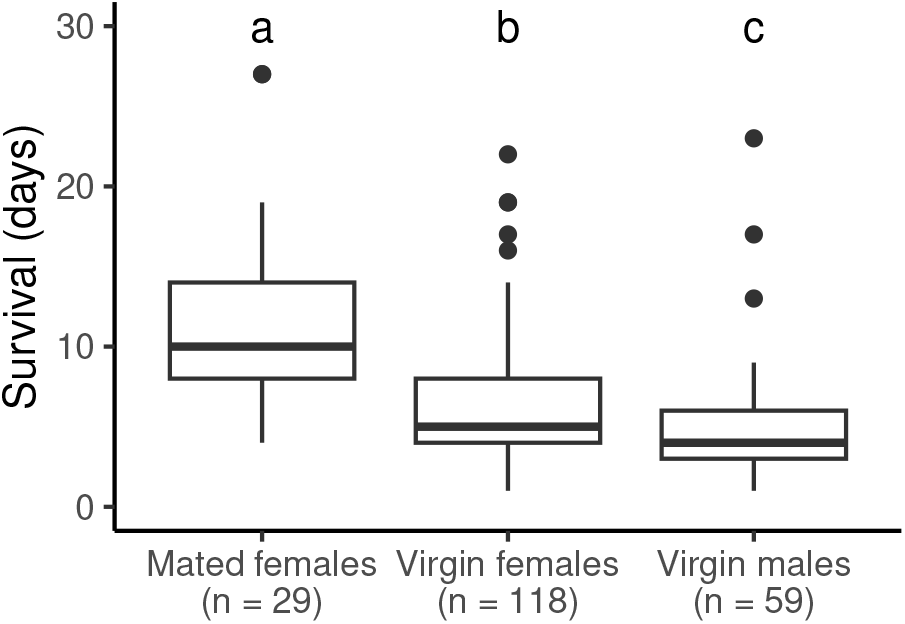
*Cydalima perspectalis* lifespan according to sex and mating status. Experiment 2017 (All males were virgin). Number of adults in each group is indicated under the group names. Significant differences between groups: global KW *p* < 0.001, pairwise differences are indicated by letters (Dunn *p* < 0.05).

Total distance flown by insects during their lifetime was tremendously variable ((0 to 153) km, *Mdn* = 10 km; Figure S3 in Sauvard et al. (2026b)). Only one insect did not fly at all during its lifetime. Total distance decreased from G1 to G3 (Figure 6A, KW *p* < 0.001), and mated females flew further than virgin ones or males (Figure 6B, KW *p* = 0.004); these tendencies were generally observed in subgroups per sex and mating status and per generation, but they were not always significant. As distance flown by mated females in a flight test did not differ from that of virgin ones or males (Figure S4 in Sauvard et al. (2026b)), the difference in flown distance seemed mainly due to their relative longer lifespan.

**Figure 6.**
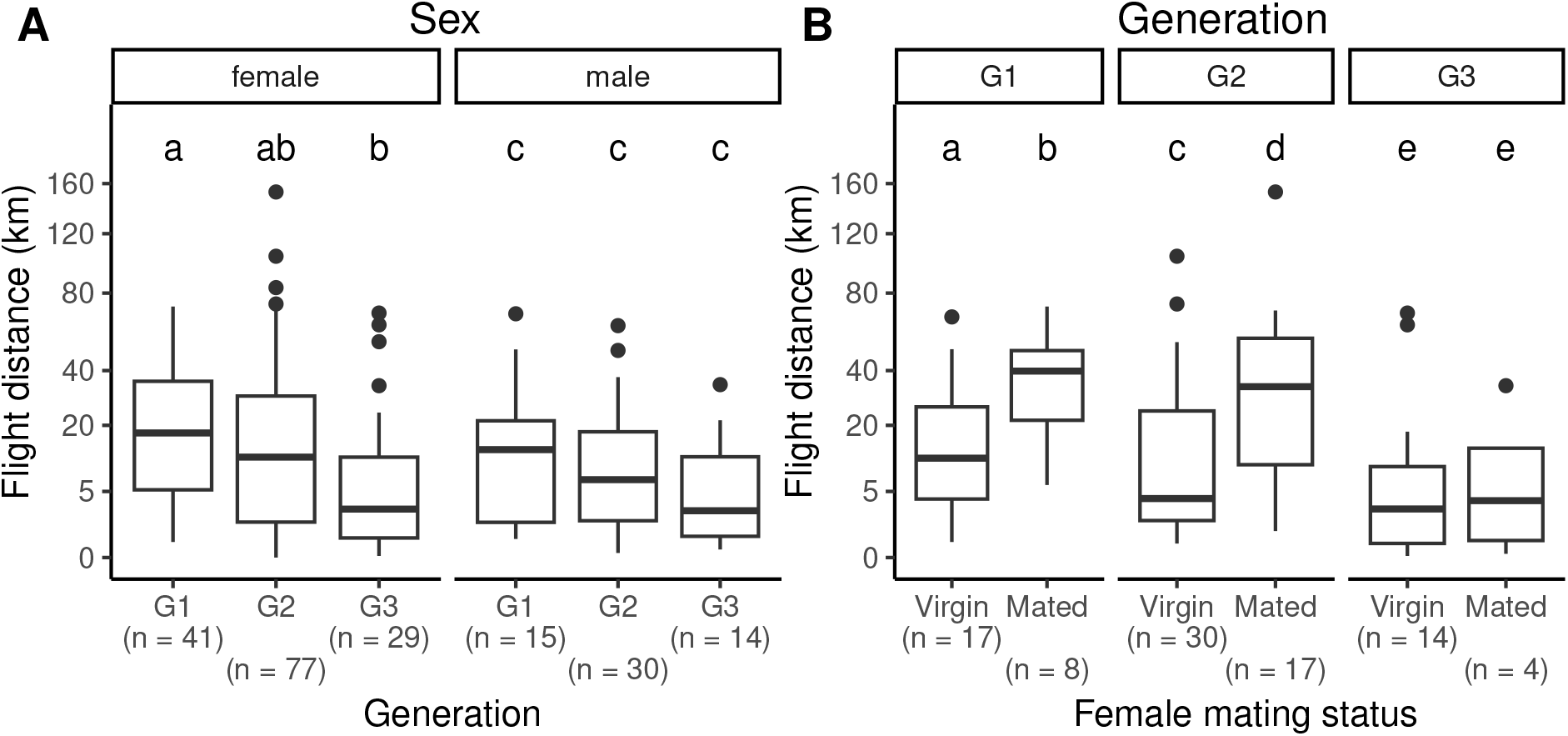
Total distances covered by *Cydalima perspectalis* adults during their entire lifetime. A. Effect of generation by sex. B. Effect of mating status by generation. Experiment 2017. Distances are displayed with square root scale to improve visibility. Number of adults in each group is indicated under the group names. Significant differences between groups: 1) global KW A female *p* = 0.004, A male NS, B G1 *p* = 0.036, B G2 *p* = 0.021, B G3 NS; 2) pairwise differences are indicated by letters (Dunn *p* < 0.05).

### Effect of flight intensity on female fitness

In 2018 supplementary experiment, female fitness was affected by flight test intensity. Their lifespan decreased with increasing flight test intensity (Figure 7; Spearman *p* < 0.001, *ρ* = *−*0.58), as did their egg laying (Figure S5 in Sauvard et al. (2026b); Spearman *p* < 0.001, *ρ* = *−*0.51). This was mainly due to a decreasing proportion of laying females (0.8, 0.67, 0.32 and 0.17 for 0, 0.25/0.5, 1 and 2/4 test nights per week, respectively, *χ*^2^ *p* = 0.002); among laying females, egg laying did not significantly decrease with increasing flight test intensity, even if the two most fecund females were both in the 0 test nights per week group.

**Figure 7.**
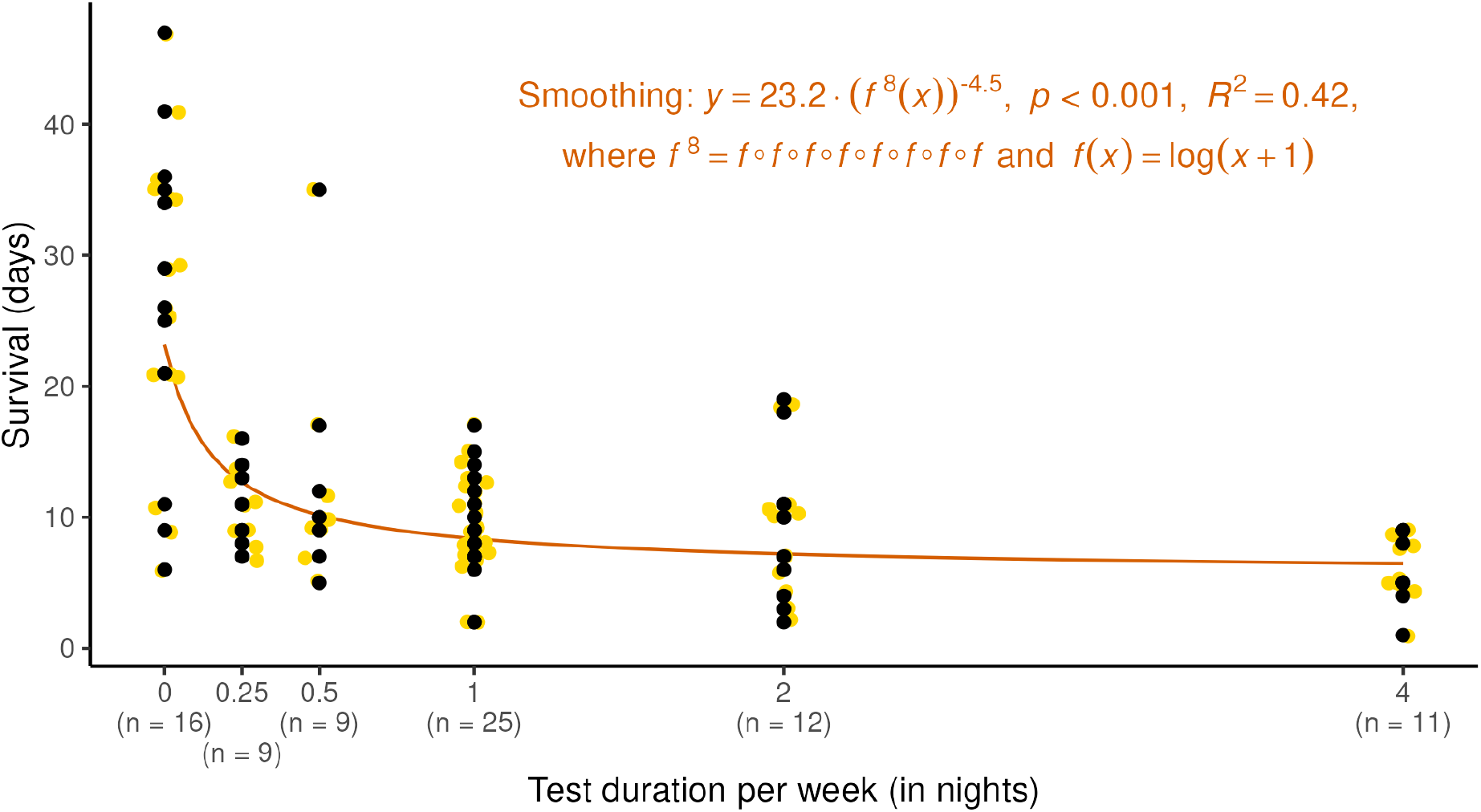
Lifespan of *Cydalima perspectalis* females under varying flight test intensity. Experiment 2018. Spearman correlation: *p* < 0.001, *ρ* = *−*0.58. Jittered gold points partially reveal the position of the overlapping points.

Even if few insects were tested in additional non-flying control protocols, their results gave interesting supplementary information on the effects of our experimental design on life-history Generation Female mating status traits. Firstly, comparison between Base and Boxwood tree controls showed that adding a Box-wood tree branch did not affect female lifespan (Figure 8A), whereas it increased their fecundity, even if the latter varied widely in both groups (Figure 8B, KW *p* = 0.025). Secondly, comparison of Tether and Fasting controls with Base control showed that female lifespan was lower in the two former protocols (Figure 8A, KW *p* = 0.007), as fecundity tended to be (Figure 8B, KW *p* = 0.07); in both cases, values in Tether and Fasting did not significantly differ. Finally, comparison of Tether and Fasting controls with corresponding flight protocol (4 test nights per week) showed that female lifespan and fecundity were slightly higher in the former protocols (Figure 8C & D), but these differences were not significant.

**Figure 8.**
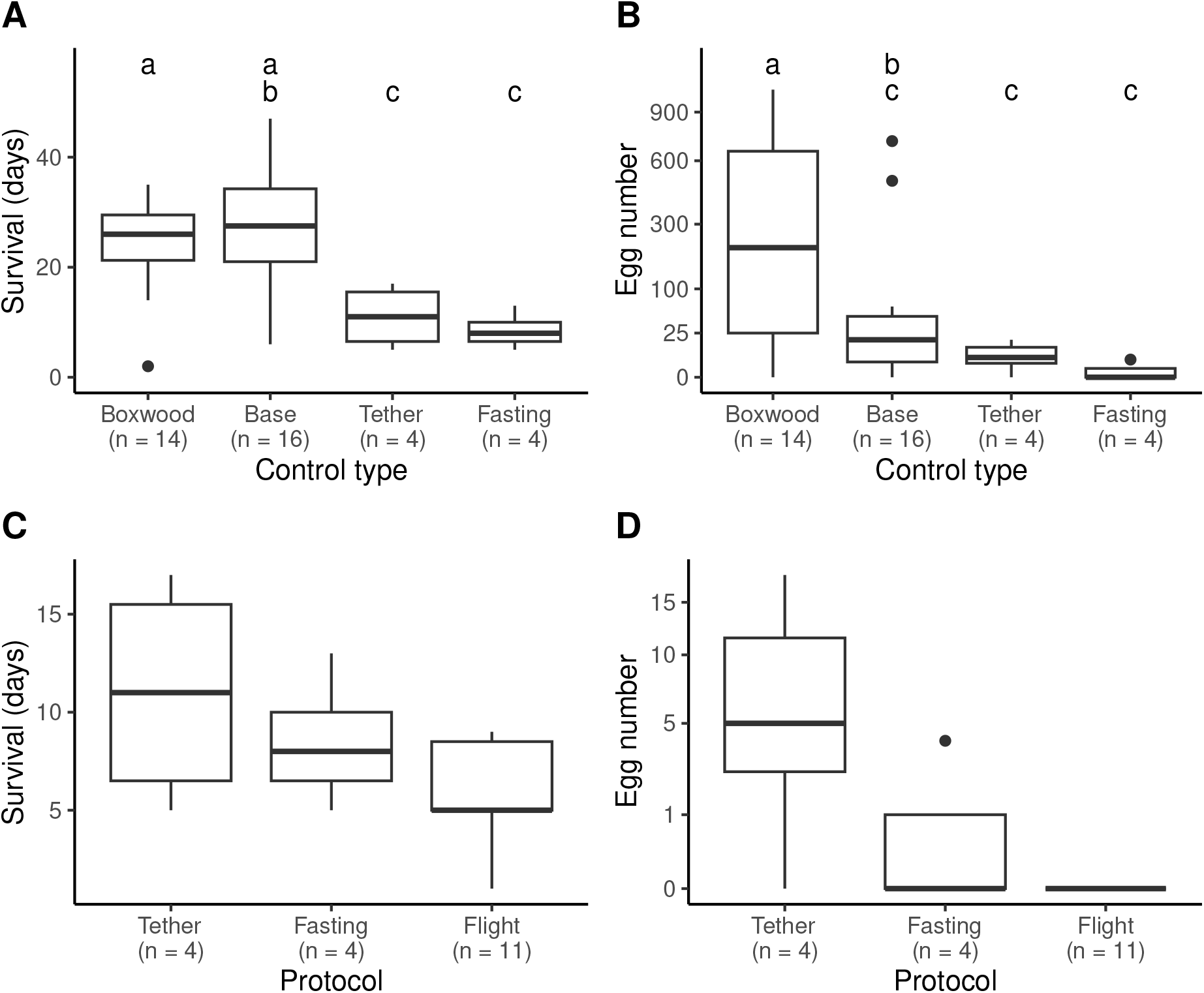
Changes in lifespan (A, C) and fecundity (B, D) for females of *Cydalima perspectalis* under varying control types (A, B), and under Tether and Fasting controls and corresponding flight protocol (4 test nights per week) (C, D). Experiment 2018. Fecundity is displayed with square root scale to improve visibility. Number of females in each group is indicated under the group names. Significant differences between groups (global KW): A, Boxwood vs Base NS, Base vs Tether vs Fasting *p* = 0.007; B, Boxwood vs Base *p* = 0.025, Base vs Tether vs Fasting *p* = 0.07; C and D: no significant differences. If appropriate, pairwise differences are indicated by letters (Dunn *p* < 0.05).

## Discussion

The box tree moth is among non-native species introduced after 2000’s and for which we have witnessed an unprecedented large and fast expansion (Bras et al., 2019; Roques et al., 2016) together with significant negative economic and ecological consequences (Ferracini et al., 2022; Kenis et al., 2013; Mitchell et al., 2018). While several studies have highlighted the role of human-activities in its rapid expansion in Europe (Bras et al., 2019, 2022), this study provides first insights of the moth’s active dispersal with the first measures of its flight capabilities. Under laboratory conditions, the tested adults were able to fly longer distances than expected, covering on average 18 km within their lifespan, with some individuals capable of performing long-distance flights (up to 150 km). The distance flown by the adults was correlated with the insects’ age and body mass, and to some extent, to the females’ mating status. We observed a trade-off between the frequency of flight tests and females’ fecundity. Finally, we found evidence for a possible generation effect, with adults from the overwintering generation presenting greater flight capabilities than the ones from the third generation. These results are a first step in understanding *C. perspectalis* natural dispersal and its role in the fast invasion in Europe. Such knowledge is required to improve and adapt current management strategies against this pest, especially in North America where it was recently accidentally introduced (Coyle et al., 2022) and populations are still localised (Seehausen et al., 2024).

### Flight performance of the box tree moth and implications in its expansion

As commonly observed with flight mill experiments in insects (Bruzzone et al., 2009; Minter et al., 2018; Pélisson et al., 2013; Sarvary et al., 2008b), we observed a considerable amount of variation in flight capabilities, but without any sex effect. Most of the *C. perspectalis* adults did short flights during a test (less than 2 km), with few individuals performing longer flights. A similar pattern was observed for the total flight distance covered over the insect’s lifespan. The majority of the adults flown about 10 km while a number of individuals were capable of covering greater distances (eg. few females flown more than 70 km). Adults able to do long-distance dispersal over their lifespan were often the ones performing a long distance flight within a flight phase. The longest flights were only performed by young adults (under 5 days-old), showing a decrease of flight capabilities with aging, which has been previously observed in other Lepidoptera (Schumacher et al., 1997; Shirai, 1998). This variable spatial dispersal ability could be due to either genetic variability or phenotypic plasticity permitting the insects to forage, search for a mate or escape unfavourable conditions. Intraspecific variation in dispersal has been suggested as an important factor for the functioning and stability of populations, allowing a species to better colonise a novel environment and be less prone to extinction (Cote et al., 2022).

Distances flown by the box tree moth during a test were comparable to distances estimated by computerised flight mills for other Lepidoptera species such as the oriental fruit moth *Cydia molesta* (Hughes and Dorn, 2002) or the smaller teo tortrix *Adoxophyes honmai* (Shirai and Kosugi, 2000). When it comes to non-native species, contrasting flight capabilities have been found depending on the studied species. Low dispersal species were often covering on average up to 2 km (e.g., emerald ash borer beetle *Agrilus planipennis*, asian long-horned beetle *Anoplophora glabripennis*, or prickly pear cactus moth *Cactoblastis cactorum* (Lopez et al., 2017; Sarvary et al., 2008a; Tussey et al., 2018)), while considered high dispersal species were covering about 7 km or more during a test (e.g., gypsy moth *Lymantria dispar*, yellow-legged hornet *Vespa velutina*, or longhorn beetle *Arhopalus rusticus* (Grilli and Fachinetti, 2018; Sauvard et al., 2018; Yang et al., 2017)). Based on the recorded flight capabilities (ie. 4.5 km on average), the box tree moth should then be considered as an intermediate dispersal species. In less than ten years, the box tree moth spread into Europe and Asia Minor and is now reported in more than 30 countries (Bras et al., 2019, 2022; Nacambo et al., 2014). Its natural dispersal was estimated at (5 to 10) km per year based on data from Germany (Casteels et al., 2011; van der Straten and Muus, 2010), while our measures on flight mills are suggesting higher dispersal capabilities. Roques et al. (2016) estimated that its invasion radius was nearly 500 km in five years. If we are taking into account the obtained flight capabilities and hypothesise that between 50 % to 25 % of the dispersing individuals is enough to establish a new population, an expansion of 10 km to 26 km per generation could be inferred. With an average of 3 generations per year in Europe, the insect would then be capable of dispersing from 30 km to 73 km a year, which is still too low to explain its fast expansion into Europe and Asia Minor. It cannot explain, for instance, the observed jumps, in a single year, between Romania (Gutue et al., 2014) and Hungary (Sáfián and Horváth, 2011) or in Spain between Galicia (Pérez-Otero et al., 2014) and Barcelona (David Castellano Franco pers. comm.), nor in France, the four new observations from 2011, which were at least 400 km apart (Brua, 2014). However, if these long distance dispersal jumps suggest a human intervention, the recorded flight capabilities could partly explain its fast expansion within some countries, such as the majority of the observed expansion between 2012 and 2016 in Croatia (Matošević et al., 2017). Similarly, surveys of adults in France in 2012 showed a colonization of approximately 100 km which is partly congruent with our findings if considering long-distance flyers. It is worth mentioning, though, that our study only tested on computerised flight mills one morph and populations of the box tree moth from one region in France. Populations can present different dispersal abilities due to local adaptation (Cote et al., 2022), while natural environmental conditions can affect species active dispersal (Cote et al., 2022; Minter et al., 2018). Moreover, Bras et al. (2019, 2022) found a significant genetic diversity within the box tree moth invasive populations in Europe. Thus, our findings would need to be confirmed by testing populations from different regions to determine potential genetic variability in flight capabilities. Recording flight performances in the field will provide complementary information on the insect’s flying behaviour and environmental factors affecting the different phases of dispersal (Cote et al., 2022). Nevertheless, these first results support the hypothesis that the box tree moth’s self-mediated dispersal together with human-mediated dispersal played a significant role in its fast expansion in Europe and Asia Minor.

### Trade-offs between flight capability, fecundity and lifespan

Dispersal is a trait affected by physical constraints which can lead to trade-offs with other life-history traits such as survival or fecundity (Bonte and Dahirel, 2017). In the box tree moth, lifespan and, in a less extent, flight capabilities over the adult lifetime, were affected by gender and mating status. Males showed the shortest lifespan, which was congruent with the observations made by Tabone et al. (2015) in other French invasive populations, and had the tendency to fly less over their lifetime compared to females. Depending on species, flight capabilities and behaviour are often sex-biased. In most of these cases, males disperse more than females, but the opposite, as in our results, was also observed in several species (Legrand et al., 2016; Turlure et al., 2011). Higher mobility in females might then contribute to a species expansion (Narimanov et al., 2022). In the box tree moth, mated females were overall living the longest and were more susceptible to sustain flight for a longer period. As a consequence, they were often covering greater distances than virgin ones during their lifespan. Contrasting observations on the influence of mating on female dispersal have been described in Lepidopteran or other insects, which suggests that it is species specific (Bloem et al., 2006; Hanski et al., 2006; Hughes and Dorn, 2002; Sarvary et al., 2008a; Shirai, 1998, 2006; Tigreros and Davidowitz, 2019). Positive tradeoff between mating and female lifespan has been reported in several insects species (Harjoko et al., 2023). Nuptial gifts from males during mating could also provide a supplement for female and potentially increase their lifespan (Wiklund et al., 1998). However, no positive effect of mating on both survival and flight capabilities has been reported to our knowledge. In the box tree moth, survival and mating likely act as a confounding factor favouring its flight capabilities. Nevertheless, assessing the influence of mating on female survival should be tested in this species to dissociate the respective contribution of each factor in favouring dispersal.

One fecunded female could be enough to establish a population as it has been advanced for the yellow-legged hornet (Arca et al., 2015; Robinet et al., 2019). Even though females of the box tree moth were mated in our flight experiments, very few eggs were found during their lifetime (less than ten) while they usually lay around 800 eggs in French invasive population (Tabone et al., 2015) and 400 eggs in the native range (Maruyama and Shinkaji, 1987; Wan et al., 2014). Our complementary experiments indicated that egg laying is greatly favoured by a stimulus from the host plant (eg. odor or texture), which can partly explain the low number of observed eggs in 2017. Moreover, the length of the tests and their repetition overtime (ie. 4 tests a week) probably exhausted the females and gave them less time to oviposit, as suggested by the lower number of eggs recorded in flying compared to non-flying adults. It has likely increased the nonoptimal conditions for oviposition and increased the confounding impacts of flight intensity and fasting on female’s fecundity, as the insects had no access to food during flight. Indeed, the fasting females in 2018 tended to have a lower fecundity than the controls while tested females on flight mills presented the lowest fecundity, even though the differences between groups were not significant. However, the harmful effect of flight on fecundity would likely be less noticeable in the field, as females have most often opportunities to feed in the night. Several studies in Lepidoptera found a negative correlation between flight duration and fecundity (Hughes and Dorn, 2002; Legrand et al., 2016; Shirai, 1995, 1998; Tigreros and Davidowitz, 2019), which could lead to egg resorption (Renault, 2020). Higher musculature in dispersing individuals together with the energy required for dispersal might then come at the expense of reproduction (Renault, 2020), which can have further impacts on species expansion. In the box tree moth, the abilities of gravid females to perform long-distance dispersal could be seen as favouring its expansion, while the potential trade-off between flight duration and fecundity would likely alleviate the possibility of establishing new populations. Yet, a dedicated study with a larger sample size and controlling for feeding behaviour (e.g. by feeding the insects, as in Sauvard et al. (2018)) would be needed to verify such a trade-off between dispersal and fecundity (Tigreros and Davidowitz, 2019).

Computerised flight mill experiments have proven to be efficient tools to estimate flight capabilities of an insect (Hughes and Dorn, 2002; Minter et al., 2018; Pélisson et al., 2013). Nevertheless, it can encourage the insects to fly longer than in the field as they do not have to support their own body mass and they are not in contact with a substrate (Bruzzone et al., 2009; Pélisson et al., 2013; Sauvard et al., 2018). On the other hand, the flight mill set up can require more energy from the insect while flying than a free flight (Minter et al., 2018): in the leafhopper *Cicadulina storeyi*, Riley et al. (1997) estimated that it requires at least (20 to 30) % more energy from the insect to overcome the friction to ‘push’ the flight mill arm than in a free flight. In our experiments, insect lifespans were rather short. While adults are supposed to live around two weeks under control conditions (Maruyama and Shinkaji, 1987; Tabone et al., 2015), only a quarter of them were alive longer than a week. Moreover, adults tested for a shorter time were overall living longer than the ones tested over a full night or for several consecutive nights. Testing the insects over a full night was likely too intensive, even with a night of break between two night tests. Furthermore, the tether glued on insect’s thorax likely affected their ability to feed themselves and thus to recover from the effort of flying on the flight mills, as it has been suggested for other insects (Le Souchu et al., 2025). This accidentally induced fastening likely decreased adults survival and could explain the decrease in body mass with aging even though insects were fed every day. These results suggests that while our experimental design in 2017 informed on the total distance the moth is physically able to cover, it likely reduced their chances to regain energy from their effort on the flight mill. Altogether, this could explain the overall shorter adult lifespan and low fecundity observed in our study.

### Seasonal variations in flight capabilities

In contradiction to our expectations, we found a seasonal variation in the flight capabilities of the box tree moth. The overwintering generation had the greatest flight capabilities. The insects were the fastest and covering the longest distances (G1; 22 km on average), while distances covered were decreasing for the second (G2; 18 km on average) and third generation (G3; 10 km on average). The insect’s pre-flight body mass presented a similar pattern, with individuals from the overwintering generation being the heaviest while the individuals from the third generation were the lightest. Insect’s mobility is often positively correlated to its body mass (Logghe et al., 2024; Minter et al., 2018). It is thus possible that the generation effect observed in the box tree moth flight capabilities primarily results of physiological variations between individuals from different generations. In the same vein, it has been observed that box tree moth overwintering generation presents higher fecundity (Wan et al., 2014) and growth rate (Leuthardt and Baur, 2013). Similarly, He et al. (2021) found that variation in photoregime was affecting the different life stages of the multivoltine fall armyworm, *Spodoptera frugiperda*, including body loss during flight.

Even though the differences between generations in the box tree moth would need to be confirmed with trials over several years, some non-exclusive hypotheses could explain the observed variations. The larval stage of the overwintering generation lasts longer than for the other generations as the insect is overwintering at the early larval instars in France (Poitou et al., 2020). Larvae have thus a longer feeding period at lower temperatures, which could explain the observed particularities for this generation. Indeed, Sarvary et al. (2008b) showed that such conditions increased the body size of the prickly pear cactus moth. In their review, Tigreros and Davidowitz (2019) found that in most Lepidoptera, food quality was negatively affecting flight performances and to a lesser extent, body size. In other words, a poor host plant quality would increase flight capabilities. Larvae of the box tree moth sequester alkaloids from their host plants whereas the amount of alkaloids in boxwood trees varies depending on the age of the leaves (Leuthardt et al., 2013). Older leaves present a higher concentration of alkaloids and Leuthardt et al. (2013) suggested that young larvae prefer them over younger leaves. Thus, the access to such resources is likely to change over the seasons. Our results suggest the last summer generation to be stationary. Therefore, females laying eggs at the end of the summer for the upcoming overwintering generation are more likely to oviposit on a plant already attacked. Taking this into consideration, we can speculate that the hatching larvae will have lower chances of having access to older leaves, as they have probably already been eaten by larvae from previous generations, and will be firstly exposed to new leaves in the spring. Feeding on a poorer host plant than the other generations, could in turn increase the flight capabilities of the overwintering generation. In addition, density-dependent dispersal can result in a decrease in flight performance at higher densities (Bonte et al., 2014; Minter et al., 2018; Renault, 2020). The generation effect in flight capabilities observed in our study could as well result from an increased population density over the generations. In the box tree moth, the second and third generations are rather stationary and can overlap, which in turn could increase local population densities (Suppo et al., 2020), and likely enhance the flight capabilities of the overwintering generation.

A combination of variation in access to resources and an increased physiological stress due to intraspecific competition may negatively affect long-distance flights (Minter et al., 2018). On the contrary, the overwintering generation might have higher flight capabilities to reduce local extinction risks due to heterogeneous habitats, inbreeding or kin competition (Bonte et al., 2014; Zilio et al., 2024). If these two hypotheses still need to be tested and the generation effects confirmed, a variation in dispersal over time can have a significant role in species management. Based on our results, the overwintering generation would be more prone to colonise new areas. This would show the importance of primarily focusing on managing or eradicating the overwintering generation. Indeed, the box tree moth is likely transported with its host plant through ornamental plant trade at the diapausing stage. Treating boxwood trees in the late summer and removing thoroughly overwintering larvae in the shrubs would not only prevent human-mediated dispersal but also limit local expansion through self-mediated dispersal in the spring.

## Conclusion

Our study provides the first findings on *C. perspectalis* flight capabilities. The box tree moth presented higher flight capabilities than expected which, however, cannot explain all the history of its invasion in Europe and Asia Minor. Moreover, the potential trade-offs between flight, fecundity and lifespan together with a seasonal variation of flight capabilities highlighted the complexity of the dispersal process in this species. Insect’s flight capabilities likely played a role in its rapid local expansion, while human-mediated dispersal favoured its long-distance dispersal. The fast spread observed within its invaded range likely results of a combination of these two processes.

## Acknowledgements

We would like to acknowledge Patrick Pineau, Philippe Lorme and Cyril Kruczkowski for helping in insect collection, Claudine Courtin and Olivier Denux for their technical assistance.

## Data, script, code, and supplementary information availability

Data and source code at the time of publication (raw log files from the flight mills, associated processed data, and R scripts) are available online in the public archive repository Recherche Data Gouv (https://doi.org/10.57745/LUWJWP; Sauvard et al. (2026a))

The Ruby pipelines used to process log files into flight data are available online in the public archive repository Recherche Data Gouv (https://doi.org/10.57745/YLMCNU; Sauvard (2025)) Supplementary material is available online in the public archive repository Recherche Data Gouv (https://doi.org/10.57745/TAN2Q5; Sauvard et al. (2026b))

## References

Arca M, Mougel F, Guillemaud T, Dupas S, Rome Q, Perrard A, Muller F, Fossoud A, Capdevielle-Dulac C, Torres-Leguizamon M, Chen XX, Tan JL, Jung C, Villemant C, Arnold G, Silvain J (2015). Reconstructing the invasion and the demographic history of the yellow-legged hornet, Vespa velutina, in Europe. Biological Invasions 17, 2357–2371. 10.1007/s10530-015-0880-9.

Bakay L, Kollár J (2018). The spread rate of Cydalima perspectalis (Walker 1859) in Slovakia (2013–2015). In: Plants and Landscape in Urban Areas. Ed. by Katarína Rovná and Ján Kollár. Slovak University of Agriculture in Nitra, Slovakia, pp. 51–54. 1015414/PUAL/2018.51-54. (Visited on 06/20/2018).

Blackburn TM, Pyšek P, Bacher S, Carlton JT, Duncan RP, Jarošík V, Wilson JRU, Richardson DM (2011). A proposed unified framework for biological invasions. Trends in Ecology & Evolution 26, 333–339. 10.1016/j.tree.2011.03.023.

Bloem S, Carpenter JE, Dorn S (2006). Mobility of mass-reared diapaused and nondiapaused Cydia pomonella (Lepidoptera : Tortricidae): Effect of mating status and treatment with gamma radiation. Journal of Economic Entomology 99, 699–706.

Bonte D, Dahirel M (2017). Dispersal: a central and independent trait in life history. Oikos 126, 472–479. 10.1111/oik.03801. (Visited on 06/27/2025).

Bonte D, De Roissart A, Wybouw N, Van Leeuwen T (2014). Fitness maximization by dispersal: evidence from an invasion experiment. Ecology 95, 3104–3111.

Bonte D, Keith S, Fronhofer EA (2024). Species interactions and eco-evolutionary dynamics of dispersal: the diversity dependence of dispersal. Philosophical Transactions of the Royal Society B: Biological Sciences 379, 20230125. 10.1098/rstb.2023.0125. (Visited on 09/17/2024).

Bowler DE, Benton TG (2005). Causes and consequences of animal dispersal strategies: relating individual behaviour to spatial dynamics. Biological Reviews 80, 205–225.10.1017/S1464793104006645.

Bras A, Auger-Rozenberg MA, Kenis M, Tabone E (2025). La pyrale du buis : un ravageur envahissant passant du jardin à la forêt. In: Invasion et expansion d’insectes bioagresseurs forestiers: Quels risques pour la forêt française dans le contexte des changements globaux ? Éditions Quæ. Collection Synthèses. Versailles, p. 309.

Bras A, Avtzis DN, Kenis M, Li H, Vétek G, Bernard A, Courtin C, Rousselet J, Roques A, Auger-Rozenberg MA (2019). A complex invasion story underlies the fast spread of the invasive box tree moth (Cydalima perspectalis) across Europe. Journal of Pest Science 92, 1187–1202. 10.1007/s10340-019-01111-x.

Bras A, Lombaert E, Kenis M, Li H, Bernard A, Rousselet J, Roques A, Auger-Rozenberg MA (2022). The fast invasion of Europe by the box tree moth: an additional example coupling multiple introduction events, bridgehead effects and admixture events. Biological Invasions. 10.1007/s10530-022-02887-3. (Visited on 08/22/2022).

Brua C (2014). La pyrale du buis:le point sur cette espèce envahissante. Phytoma 675, 16–22.

Bruzzone OA, Villacide JM, Bernstein C, Corley JC (2009). Flight variability in the woodwasp Sirex noctilio (Hymenoptera: Siricidae): an analysis of flight data using wavelets. Journal of Experimental Biology 212, 731–737. 10.1242/jeb.022517.

Casteels H, Witters J, Vandierendonck S, Van Remoortere L, Goossens F (2011). First report of Cydalima perspectalis (Lepidoptera: Crambidae) in Belgium. In: Proceedings of the 63rd International Symposium on Crop Protection, Ghent, pp. 151–155.

Coombs M (1997). Tethered-flight and age-related reproductive performance of Helicoverpa punctigera (Wallengren) and H. armigera (Hubner) (Lepidoptera: Noctuidae). Australian Journal of Zoology 45, 409–422. 10.1071/ZO96064.

Cote J, Brodin T, Fogarty S, Sih A (2017). Non-random dispersal mediates invader impacts on the invertebrate community. Journal of Animal Ecology 86, 1298–1307.10.1111/1365-2656.12734. (Visited on 06/27/2025).

Cote J, Dahirel M, Schtickzelle N, Altermatt F, Ansart A, Blanchet S, Chaine AS, De Laender F, De Raedt J, Haegeman B, Jacob S, Kaltz O, Laurent E, Little CJ, Madec L, Manzi F, Masier S, Pellerin F, Pennekamp F, Therry L, et al. (2022). Dispersal syndromes in challenging environments: A cross-species experiment. Ecology Letters, ele.14124. 10.1111/ele.14124. (Visited on 11/23/2022).

Coyle DR, Adams J, Bullas-Appleton E, Llewellyn J, Rimmer A, Skvarla MJ, Smith SM, Chong JH (2022). Identification and Management of Cydalima perspectalis (Lepidoptera: Crambidae) in North America. Journal of Integrated Pest Management 13. Ed. by Carlos Bogran, 24. 10.1093/jipm/pmac020. (Visited on 09/22/2022).

Davis AK, Chi J, Bradley C, Altizer S (2012). The Redder the Better: Wing Color Predicts Flight Performance in Monarch Butterflies. Plos One 7. 10.1371/journal.pone.0041323.

Di Domenico F, Lucchese F, Magri D (2012). Buxus in Europe: Late Quaternary dynamics and modern vulnerability. Perspectives in Plant Ecology Evolution and Systematics 14, 354–362. 10.1016/j.ppees.2012.07.001.

EPPO (2012). EPPO Technical Document No. 1061, EPPO Study on the Risk of Imports of Plants for Planting. EPPO Paris. 75 pp.

Essl F, Bacher S, Genovesi P, Hulme PE, Jeschke JM, Katsanevakis S, Kowarik I, Kühn I, Pyšek P, Rabitsch W, Schindler S, van Kleunen M, Vilà M, Wilson JRU, Richardson DM (2018). Which Taxa Are Alien? Criteria, Applications, and Uncertainties. Bioscience 68, 496–509. 10.1093/biosci/biy057.

Estoup A, Guillemaud T (2010). Reconstructing routes of invasion using genetic data: why, how and so what? Molecular Ecology 19, 4113–4130. 10.1111/j.1365-294X.2010.04773.x.

Fenn-Moltu G, Ollier S, Caton B, Liebhold AM, Nahrung H, Pureswaran DS, Turner RM, Yamanaka T, Bertelsmeier C (2022). Alien insect dispersal mediated by the global movement of commodities. Ecological Applications. 10.1002/eap.2721. (Visited on 08/29/2022).

Ferracini C, Pogolotti C, Mancardi P, Miglio M, Bonelli S, Barbero F (2022). The Box Tree Moth: An Invasive Species Severely Threatening Buxus Natural Formation in NW Italy. Forests 13, 178. 10.3390/f13020178. (Visited on 01/27/2022).

Garnas JR, Auger-Rozenberg MA, Roques A, Bertelsmeier C, Wingfield MJ, Saccaggi DL, Roy HE, Slippers B (2016). Complex patterns of global spread in invasive insects: eco-evolutionary and management consequences. Biological Invasions 18, 935–952. 10.1007/s10530-016-1082-9.

Gil-Vives L, Compa M, Sureda A, Pinya S (2026). Impacts of the Invasive Moth Cydalima perspectalis on the Native Shrub Buxus balearica in Mallorca (Spain). Chemistry & Biodiversity 23, e02516. 10.1002/cbdv.202502516. (Visited on 03/16/2026).

Gninenko YI, Shiryaeva NV, Shurov VI (2014). The box tree moth-a new invasive pest in the Caucasian Forests. Plant Health -Research and Practice 1, 32–39.

Göttig S, Herz A (2017). Observations on the seasonal flight activity of the box tree pyralid Cydalima perspectalis (Lepidoptera: Crambidae) in the Rhine-Main Region of Hessia. Journal für Kulturpflanzen 69, 157–165. 10.1399/JfK.2017.05.01.

Grilli MP, Fachinetti R (2018). The role of host patch characteristics and dispersal capability in distribution and abundance of Arhopalus rusticus in central Argentina. Entomologia Experimentalis et Applicata 166, 183–190. 10.1111/eea.12653.

Gutue C, Gutue M,, Rosca I (2014). Crambidae associated with parks and ornamental gardens of Bucharest. Scientific Papers. Series B, Horticulture LVIII, 323–326.

Hanski I, Saastamoinen M, Ovaskainen O (2006). Dispersal-related life-history trade-offs in a butterfly metapopulation. Journal of Animal Ecology 75, 91–100. 10.1111/j.1365-2656.2005.01024.x. (Visited on 08/01/2018).

Harjoko DN, Hua QQH, Toh EMC, Goh CYJ, Puniamoorthy N (2023). A window into fly sex: mating increases female but reduces male longevity in black soldier flies. Animal Behaviour 200, 25–36. 10.1016/j.anbehav.2023.03.007. (Visited on 02/12/2026).

He L, Ge S, Zhang H, He W, Yan R, Wu K (2021). Photoregime Affects Development, Reproduction, and Flight Performance of the Invasive Fall Armyworm (Lepidoptera: Noctuidae) in China. Environmental Entomology 50, 367–381. 10.1093/ee/nvaa172. (Visited on 08/08/2024).

Hoddle MS, Hoddle CD, Faleiro JR, El-Shafie HAF, Jeske DR, Sallam AA (2015). How Far Can the Red Palm Weevil (Coleoptera: Curculionidae) Fly?: Computerized Flight Mill Studies With Field-Captured Weevils. Journal Of Economic Entomology 108, 2599–2609. 10.1093/jee/tov240.

Hughes J, Dorn S (2002). Sexual differences in the flight performance of the oriental fruit moth, Cydia molesta. Entomologia Experimentalis et Applicata 103, 171–182. 10.1046/j.1570-7458.2002.00967.x.

Hulme PE (2009). Trade, transport and trouble: managing invasive species pathways in an era of globalization. Journal of Applied Ecology 46, 10–18. 10.1111/j.13652664.2008.01600.x.

Kawazu K, Nakamura S, Adati T (2010a). Rearing of the box tree pyralid, Glyphodes perspectalis, larvae using an artificial diet. Applied Entomology and Zoology 45, 163–168. 10.1303/aez.2010.163.

Kawazu K, Nakamura S, Honda H, Adati T (2010b). Effects of photoregime on the diel rhythmicity of male responses to sex pheromones in Glyphodes perspectalis (Lepidoptera: Crambidae).Applied Entomology and Zoology 45, 169–176. 10.1303/aez.2010.169.

Kazilas C, Kalaentzis K, Demetriou J, Koutsoukos E, Strachinis I, Andriopoulos P (2021). Utilization of citizen science data to monitor alien species: the box tree moth Cydalima perspectalis (Walker, 1859) (Lepidoptera: Crambidae) invades natural vegetation in Greece. BioInvasions Records 10, 1032–1044. 10.3391/bir.2021.10.4.28. (Visited on 05/24/2022).

Kenis M, Nacambo S, Leuthardt FLG, Di Domenico F, Haye T (2013). The box tree moth, Cydalima perspectalis, in Europe: horticultural pest or environmental disaster? Aliens 33, 38–41.

Krüger EO (2008). Glyphodes perspectalis (Walker, 1859) -Neu für die Fauna Europas (Lepidoptera: Crambidae). Entomologische Zeitschrift mit Insekten-Börse 118, 81–83.

Le Souchu E, Sallé A, Bankhead-Dronnet S, Laparie M, Sauvard D (2025). Intra-and interspecific variations in flight performance of oak-associated Agrilinae (Coleoptera: Buprestidae) using computerised flight mills. Peer Community Journal 5, e64. 10.24072/pcjournal.560. (Visited on 09/05/2025).

Ledru L, Garnier J, Gallet C, Noûs C, Ibanez S (2022). Spatial structure of natural boxwood and the invasive box tree moth can promote coexistence. Ecological Modelling 465, 109844. 10.1016/j.ecolmodel.2021.109844. (Visited on 04/08/2022).

Legrand D, Larranaga N, Bertrand R, Ducatez S, Calvez O, Stevens VM, Baguette M (2016). Evolution of a butterfly dispersal syndrome. Proceedings of the Royal Society B: Biological Sciences 283, 20161533. 10.1098/rspb.2016.1533. (Visited on 01/23/2026).

Leuthardt FLG, Baur B (2013). Oviposition preference and larval development of the invasive moth Cydalima perspectalis on five European box-tree varieties. Journal of Applied Entomology 137, 437–444. 10.1111/jen.12013.

Leuthardt FLG, Glauser G, Baur B (2013). Composition of alkaloids in different box tree varieties and their uptake by the box tree moth Cydalima perspectalis. Chemoecology 23, 203–212. 10.1007/s00049-013-0134-1. (Visited on 10/18/2014).

Logghe G, Taelman C, Van Hecke F, Batsleer F, Maes D, Bonte D (2024). Unravelling arthropod movement in natural landscapes: Small-scale effects of body size and weather conditions. Journal of Animal Ecology 93, 1365–1379. 10.1111/1365-2656.14161. (Visited on 10/26/2024).

Lopez VM, Hoddle MS, Francese JA, Lance DR, Ray AM (2017). Assessing Flight Potential of the Invasive Asian Longhorned Beetle (Coleoptera: Cerambycidae) With Computerized Flight Mills. Journal of Economic Entomology 110, 1070–1077. 10.1093/jee/tox046.

Maruyama T, Shinkaji N (1987). Studies on the life-cycle of the box-tree pyralid, Glyphodes perspectalis (Walker) (Lepidoptera: Pyralidae). I. Seasonal adult emergence and developmental velocity. Japanese Journal of Applied Entomology and Zoology 31, 226–232.

Maruyama T, Shinkaji N (1991). The life-cycle of the box-tree pyralid, Glyphodes perspectalis (Walker) (Lepidoptera: Pyralidae). II. Developmental characteristics of larvae. Japanese Journal of Applied Entomology and Zoology 35, 221–230.

Matošević D, Lukić I, Bras A, Lacković N, Pernek M (2017). Spatial Distribution, Genetic Diversity and Food Choice of Box Tree Moth (Cydalima perspectalis) in Croatia. SEEFOR–South-East European Forestry 8, 41–46. 10.15177/seefor.17-06.

Minter M, Pearson A, Lim KS, Wilson K, Chapman JW, Jones CM (2018). The tethered flight technique as a tool for studying life-history strategies associated with migration in insects. Ecological Entomology 43, 397–411. 10.1111/een.12521. (Visited on 07/16/2018).

Mitchell R, Chitanava S, Dbar R, Kramarets V, Lehtijärvi A, Matchutadze I, Mamadashvili G, Matsiakh I, Nacambo S, Papazova-Anakieva I, Sathyapala S, Tuniyev B, Vétek G, Zukhbaia M, Kenis M (2018). Identifying the ecological and societal consequences of a decline in Buxus forests in Europe and the Caucasus. Biological Invasions 20, 3605–3620. 10.1007/s10530-018-1799-8.

Nacambo S, Leuthardt FLG, Wan H, Li H, Haye T, Baur B, Weiss RM, Kenis M (2014). Development characteristics of the box-tree moth Cydalima perspectalis and its potential distribution in Europe. Journal of Applied Entomology 138, 14–26. 10.1111/jen.12078.

Narimanov N, Bauer T, Bonte D, Fahse L, Entling MH (2022). Accelerated invasion through the evolution of dispersal behaviour. Global Ecology and Biogeography 31, 2423–2436. 10.1111/geb.13599. (Visited on 11/23/2022).

Pélisson PF, Bernstein C, Débias F, Menu F, Venner S (2013). Dispersal and dormancy strategies among insect species competing for a pulsed resource. Ecological Entomology 38, 470–477. 10.1111/een.12038.

Pérez-Otero R, Mansilla JP, Vidal M (2014). Cydalima perspectalis Walker, 1859 (Lepidoptera, Crambidae): una nueva amenaza para Buxus spp. en la Península Ibérica. Arquivos Entomolóx-icos 10, 225–228.

Perrings C, Dehnen-Schmutz K, Touza J, Williamson M (2005). How to manage biological invasions under globalization. Trends in Ecology & Evolution 20, 212–215.10.1016/j.tree.2005.02.011.

Poitou L, Bras A, Pineau P, Lorme P, Roques A, Rousselet J, Auger-Rozenberg MA, Laparie M (2020). Diapause Regulation in Newly Invaded Environments: Termination Timing Allows Matching Novel Climatic Constraints in the Box Tree Moth, Cydalima perspectalis (Lepidoptera:Crambidae). Insects 11, 629. 10.3390/insects11090629. (Visited on 09/14/2020).

Qin J, Liu Y, Zhang L, Cheng Y, Sappington TW, Jiang X (2018). Effects of Moth Age and Rearing Temperature on the Flight Performance of the Loreyi Leafworm, Mythimna loreyi (Lepidoptera: Noctuidae), in Tethered and Free Flight. Journal of Economic Entomology 111, 1243–1248. 10.1093/jee/toy076.

R Core Team (2022). R: A Language and Environment for Statistical Computing. Version 4.2.2. R Foundation for Statistical Computing. Vienna, Austria. URL: https://www.R-project.org/.

Renault D (2020). A Review of the Phenotypic Traits Associated with Insect Dispersal Polymorphism, and Experimental Designs for Sorting out Resident and Disperser Phenotypes. Insects 11, 214. 10.3390/insects11040214. (Visited on 01/22/2026).

Renault D, Laparie M, McCauley SJ, Bonte D (2018). Environmental Adaptations, Ecological Filtering, and Dispersal Central to Insect Invasions. Annual Review of Entomology 63, 345–368. 10.1146/annurev-ento-020117-043315.

Renault D, Rantier Y, Convey P, Bergerot B (2025). Evolution of dispersal capacities during range expansion: size and behaviour matter in an arthropod invading the sub-Antarctic Kerguelen archipelago. Proceedings of the Royal Society B: Biological Sciences 292, 20251136. 10.1098/rspb.2025.1136. (Visited on 01/22/2026).

Riley JR, Downham MCA, Cooter RJ (1997). Comparison of the performance of Cicadulina leafhoppers on flight mills with that to be expected in free flight. Entomologia Experimentalis et Applicata 83, 317–322. 10.1046/j.1570-7458.1997.00186.x. (Visited on 07/25/2018).

Robinet C, Darrouzet É, Suppo C (2019). Spread modelling: a suitable tool to explore the role of human-mediated dispersal in the range expansion of the yellow-legged hornet in Europe. International Journal of Pest Management 65, 258–267. 10.1080/09670874.2018.1484529.

Robinet C, Imbert CÉ, Rousselet J, Sauvard D, Garcia J, Goussard F, Roques A (2012). Human-mediated long-distance jumps of the pine processionary moth in Europe. Biological Invasions 14, 1557–1569. 10.1007/s10530-011-9979-9.

Robinet C, Suppo C, Darrouzet É (2017). Rapid spread of the invasive yellow-legged hornet in France: the role of human-mediated dispersal and the effects of control measures. Journal of Applied Ecology 54, 205–215. 10.1111/1365-2664.12724.

Roques A, Auger-Rozenberg MA, Blackburn TM, Garnas J, Pyšek P, Rabitsch W, Richardson DM, Wingfield MJ, Liebhold AM, Duncan RP (2016). Temporal and interspecific variation in rates of spread for insect species invading Europe during the last 200 years. Biological Invasions 18, 907–920. 10.1007/s10530-016-1080-y.

Sáfián S, Horváth B (2011). Box Tree Moth – Cydalima perspectalis (Walker, 1859), new member in the Lepidoptera fauna of Hungary (Lepidoptera: Crambidae). Natura Somogyiensis 19, 245–246.

Sakai AK, Allendorf FW, Holt JS, Lodge DM, Molofsky J, With KA, Baughman S, Cabin RJ, Cohen JE, Ellstrand NC, McCauley DE, O’Neil P, Parker IM, Thompson JN, Weller SG (2001). The population biology of invasive species. Annual Review of Ecology And Systematics 32, 305–332. 10.1146/annurev.ecolsys.32.081501.114037.

Sarvary MA, Bloem KA, Bloem S, Carpenter JE, Hight SD, Dorn S (2008a). Diel Flight Pattern and Flight Performance of Cactoblastis cactorum (Lepidoptera: Pyralidae) Measured on a Flight Mill: Influence of Age, Gender, Mating Status, and Body Size. Journal of Economic Entomology 101, 314–324. 10.1093/jee/101.2.314.

Sarvary MA, Hight SD, Carpenter JE, Bloem S, Bloem KA, Dorn S (2008b). Identification of Factors Influencing Flight Performance of Field-Collected and Laboratory-Reared, Overwintered, and Nonoverwintered Cactus Moths Fed with Field-Collected Host Plants. Environmental Entomology 37, 1291–1299. 10.1093/ee/37.5.1291.

Sauvard D (2025). Scripts to manage log files from URZF flight mills (2024). Version V1. 10.57745/YLMCNU.

Sauvard D, Bras A, Auger-Rozenberg MA, Rousselet J, Roques A (2026a). Flight performance of the highly invasive box tree moth Cydalima perspectalis (Lepidoptera: Crambidae). Version DRAFT VERSION. 10.57745/LUWJWP.

Sauvard D, Bras A, Auger-Rozenberg MA, Rousselet J, Roques A (2026b). Supplement for article Flight performance of the highly invasive box tree moth Cydalima perspectalis (Lepidoptera: Crambidae). Version DRAFT VERSION. 10.57745/TAN2Q5.

Sauvard D, Imbault V, Darrouzet É (2018). Flight capacities of yellow-legged hornet (Vespa velutina nigrithorax, Hymenoptera: Vespidae) workers from an invasive population in Europe. Plos One 13. 10.1371/journal.pone.0198597.

Schmera D, Baur B (2024). Climate Warming and the Dynamics of the Invasive Box-Tree Moth Cydalima perspectalis in the Suburbs of Basel (Switzerland) and in the Nearby Natural Box-Tree Forest: A 15-Year Study. Journal of Applied Entomology n/a, 1–10. 10.1111/jen.13381. (Visited on 01/07/2025).

Schumacher P, Weyeneth A, Weber DC, Dorn S (1997). Long flights in Cydia pomonella L. (Lepidoptera: Tortricidae) measured by a flight mill: influence of sex, mated status and age. Physiological Entomology 22, 149–160. 10.1111/j.1365-3032.1997.tb01152.x.

Seebens H, Blackburn TM, Dyer EE, Genovesi P, Hulme PE, Jeschke JM, Pagad S, Pyšek P, Winter M, Arianoutsou M, Bacher S, Blasius B, Brundu G, Capinha C, Celesti-Grapow L, Dawson W, Dullinger S, Fuentes N, Jäger H, Kartesz J, et al. (2017). No saturation in the accumulation of alien species worldwide. Nature Communications 8. 10.1038/ncomms14435.

Seehausen ML, Rimmer A, Wiesner A, Kenis M, Scott-Dupree C, Smith SM (2024). Modelling potential distribution of the invasive box tree moth across Asia, Europe, and North America. Plos One 19, e0302259. 10.1371/journal.pone.0302259. (Visited on 04/29/2024).

Shirai Y (1995). Longevity, flight ability and reproductive performance of the diamondback moth, Plutella xylostella (L) (Lepidoptera: Yponomeutidae), related to adult body size. Researches on Population Ecology 37, 269–277. 10.1007/BF02515829.

Shirai Y (1998). Laboratory evaluation of flight ability of the Oriental corn borer, Ostrinia furnacalis (Lepidoptera : Pyralidae). Bulletin of Entomological Research 88, 327–333. 10.1017/S0007485300025943.

Shirai Y (2006). Flight activity, reproduction, and adult nutrition of the beet webworm, Spoladea recurvalis (Lepidoptera : Pyralidae). Applied Entomology and Zoology 41, 405–414. 10.1303/aez.2006.405.

Shirai Y, Kosugi Y (2000). Flight activity of the smaller tea tortrix, Adoxophyes honmai (Lepidoptera : Tortricidae). Applied Entomology and Zoology 35, 459–466. 10.1303/aez.2000.459.

Simberloff D, Martin JL, Genovesi P, Maris V, Wardle DA, Aronson J, Courchamp F, Galil B, García-Berthou E, Pascal M, Pyšek P, Sousa R, Tabacchi É, Vilà M (2013). Impacts of biological invasions: what’s what and the way forward. Trends in Ecology & Evolution 28, 58–66. 10.1016/j.tree.2012.07.013.

Suppo C, Bras A, Robinet C (2020). A temperature-and photoperiod-driven model reveals complex temporal population dynamics of the invasive box tree moth in Europe. Ecological Modelling 432, 109229. 10.1016/j.ecolmodel.2020.109229. (Visited on 08/03/2020).

Tabone É, Enriquez T, Giorgi C, Venard M, Colombel E, Gaglio F, Buradino M, Martin JC (2015). Mieux connaître la pyrale du buis Cydalima perspectalis. Phytoma 685, 18–20. URL: https://hal.inrae.fr/hal-02634876.

Tigreros N, Davidowitz G (2019). Flight-fecundity tradeoffs in wing-monomorphic insects. In: Advances in Insect Physiology. Vol. 56. Academic Press, pp. 1–41. 10.1016/bs.aiip.2019.02.001. (Visited on 02/06/2026).

Turlure C, Baguette M, Stevens VM, Maes D (2011). Species-and sex-specific adjustments of movement behavior to landscape heterogeneity in butterflies. Behavioral Ecology 22, 967–975. 10.1093/beheco/arr077. (Visited on 06/18/2025).

Tussey DA, Aukema BH, Charvoz AM, Venette RC (2018). Effects of Adult Feeding and Overwintering Conditions on Energy Reserves and Flight Performance of Emerald Ash Borer (Coleoptera: Buprestidae). Environmental Entomology 47, 755–763. 10.1093/ee/nvy040.

van der Straten MJ, Muus TST (2010). The box tree pyralid, Glyphodes perspectalis (Lepidoptera: Crambidae), an invasive alien moth ruining box trees. Proceedings of the Netherlands Entomological Society Meeting 21, 107–111.

Wan H, Haye T, Kenis M, Nacambo S, Xu H, Zhang F, Li H (2014). Biology and natural enemies of Cydalima perspectalis in Asia: Is there biological control potential in Europe? Journal of Applied Entomology 138, 715–722. 10.1111/jen.12132.

Wiklund C, Kaitala A, Wedell N (1998). Decoupling of reproductive rates and parental expenditure in a polyandrous butterfly. Behavioral Ecology 9, 20–25. 10.1093/beheco/9.1.20. (Visited on 02/12/2026).

Wilson JRU, Dormontt EE, Prentis PJ, Lowe AJ, Richardson DM (2009). Something in the way you move: dispersal pathways affect invasion success. Trends in Ecology & Evolution 24, 136–144. 10.1016/j.tree.2008.10.007.

Yang F, Luo Y, Shi J (2017). The influence of geographic population, age, and mating status on the flight activity of the Asian gypsy moth Lymantria dispar (Lepidoptera: Erebidae) in China. Applied Entomology and Zoology 52, 265–270. 10.1007/s13355-016-0475-7.

Zilio G, Deshpande JN, Duncan AB, Fronhofer EA, Kaltz O (2024). Dispersal evolution and eco-evolutionary dynamics in antagonistic species interactions. Trends in Ecology & Evolution 0. 10.1016/j.tree.2024.03.006. (Visited on 04/22/2024).

