## Supplementary Material for "Flight performance of the highly invasive box tree moth *Cydalima perspectalis* (Lepidoptera: Crambidae)"

**Table S1** – Statistics for flight parameters for *Cydalima perspectalis* for each generation and sex. Experiment 2017, only first test of each insect. The modalities A-Normal and B-Intensive were mixed as no difference was observed. The column Generation effect gives *p*-values of between generation KW tests; if appropriate, pairwise differences are indicated below by letters (G1 G2 G3; Dunn $p < 0.05$). Stars, if any, indicate a significant difference between sexes (KW, *: $p < 0.05$, ***: $p < 0.001$).

| | | G1 | | G2 | | | G3 | | | Generation effect | |
|---|---|---|---|---|---|---|---|---|---|---|---|
| | | F | M | F | | M | F | | M | F | M |
| Total distance (km) | Mdn | 5.592 | 8.823 | 1.025 | | 2.586 | 0.529 | | 1.220 | < 0.001 (a b c) | 0.024 (a ab b) |
| | Q3 | 13.163 | 13.197 | 5.489 | | 9.340 | 1.500 | | 2.538 | | |
| | Max | 29.549 | 22.718 | 34.666 | | 36.105 | 17.443 | | 6.826 | | |
| Total flight duration (h) | Mdn | 2.4 | 3.6 | 0.5 | | 1.0 | 0.3 | | 0.7 | 0.001 (a b b) | 0.059 |
| | Q3 | 4.4 | 4.2 | 2.2 | | 3.2 | 0.8 | | 1.5 | | |
| | Max | 8.1 | 6.8 | 12.6 | | 9.8 | 7.5 | | 3.1 | | |
| Mean flight phase distance (km) | Mdn | 0.058 | 0.061 | 0.024 | | 0.049 | 0.011 | | 0.023 | 0.002 (a a b) | 0.247 |
| | Q3 | 0.196 | 0.214 | 0.127 | | 0.160 | 0.044 | | 0.056 | | |
| | Max | 2.115 | 0.575 | 0.993 | | 1.590 | 0.249 | | 0.490 | | |
| Max flight phase distance (km) | Mdn | 0.976 | 1.658 | 0.267 | | 0.698 | 0.074 | | 0.280 | < 0.001 (a b c) | 0.068 |
| | Q3 | 2.997 | 3.288 | 1.519 | | 2.391 | 0.316 | | 0.359 | | |
| | Max | 21.546 | 6.770 | 11.648 | | 13.331 | 10.121 | | 5.108 | | |
| Mean speed (m s$^{-1}$) | M | 0.667 | 0.625 | 0.622 | | 0.656 | 0.474 | | 0.496 | < 0.001 (a a b) | 0.007 (ab a b) |
| | SD | 0.176 | 0.182 | 0.167 | | 0.169 | 0.135 | | 0.090 | | |
| | Max | 1.045 | 0.927 | 1.055 | | 0.981 | 0.718 | | 0.645 | | |
| Max speed (m s$^{-1}$) | M | 0.967 | 0.922 | 0.854 | | 0.890 | 0.661 | | 0.629 | < 0.001 (a b c) | < 0.001 (a a b) |
| | SD | 0.237 | 0.223 | 0.220 | | 0.221 | 0.238 | | 0.104 | | |
| Mass (g) | Mdn | 0.093 | 0.090 | 0.098 | | 0.086 | 0.077 | | 0.076 | < 0.001 (a a b) | 0.091 |
| | Q3 | 0.113 | 0.101 | 0.109 | *** | 0.090 | 0.083 | | 0.094 | | |
| | Max | 0.185 | 0.117 | 0.139 | | 0.119 | 0.132 | | 0.108 | | |
| Mass loss (g) | Mdn | 0.014 | 0.016 | 0.013 | | 0.015 | 0.011 | | 0.013 | < 0.001 (a a b) | 0.034 (a ab b) |
| | Q3 | 0.020 | 0.019 | 0.016 | * | 0.018 | 0.012 | | 0.016 | | |
| | Max | 0.027 | 0.028 | 0.025 | | 0.021 | 0.026 | | 0.019 | | |
| Relative mass loss | Mdn | 0.170 | 0.193 | 0.136 | | 0.172 | 0.131 | | 0.174 | 0.005 (a b b) | 0.239 |
| | Q3 | 0.196 | 0.217 | 0.157 | *** | 0.191 | 0.166 | * | 0.193 | | |
| | Max | 0.325 | 0.252 | 0.264 | | 0.242 | 0.198 | | 0.219 | | |

G1, G2, G3: first, second, third generation; F, M: females, males.

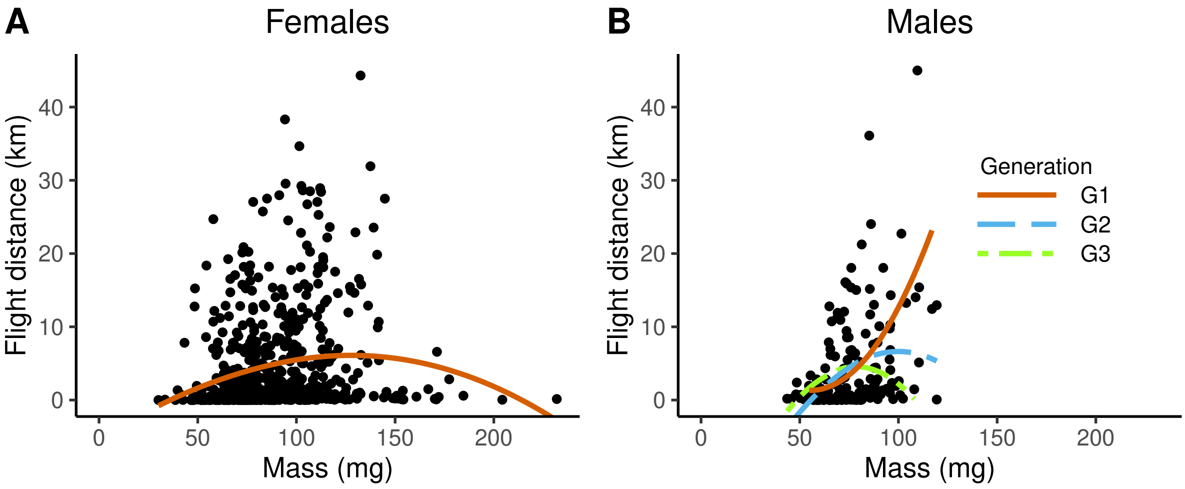


**Figure S1** – Total distances covered by *Cydalima perspectalis* females (A) and males (B) during a flight test according to their mass at the beginning of the test. Experiment 2017 (All adults gathered). The colored lines are loess smoothings of dot clouds (All females merged, males separated by generation).

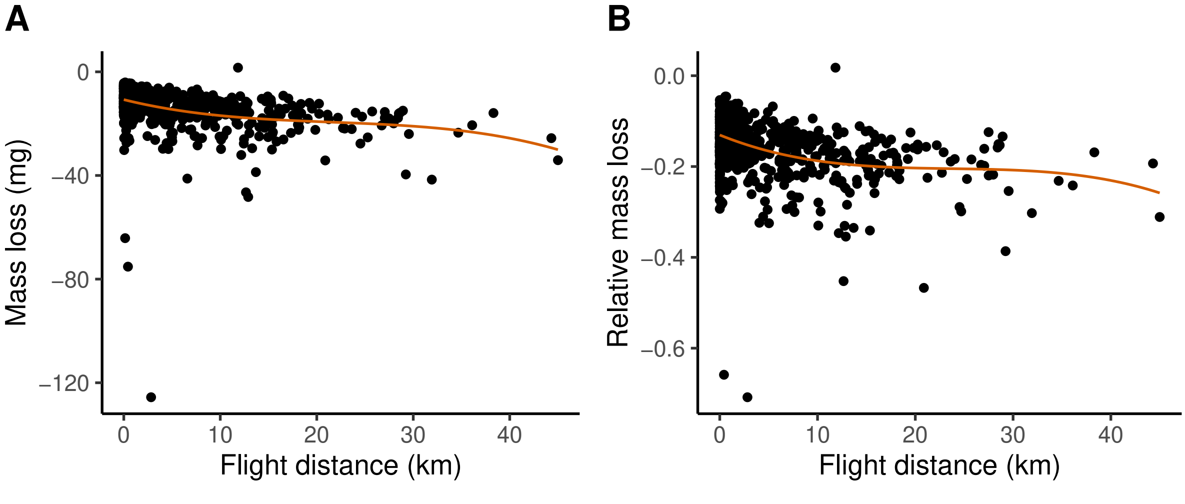


**Figure S2** – Changes in *Cydalima perspectalis* mass loss (A) and relative mass loss (B) during a flight test according to the total distance the insect covered during the test. Experiment 2017 (All adults gathered). Global correlations (Spearman): A $p < 0.001$ $\rho = -0.48$, B $p < 0.001$ $\rho = -0.38$. The red lines are degree 3 polynomial smoothings of dot clouds.

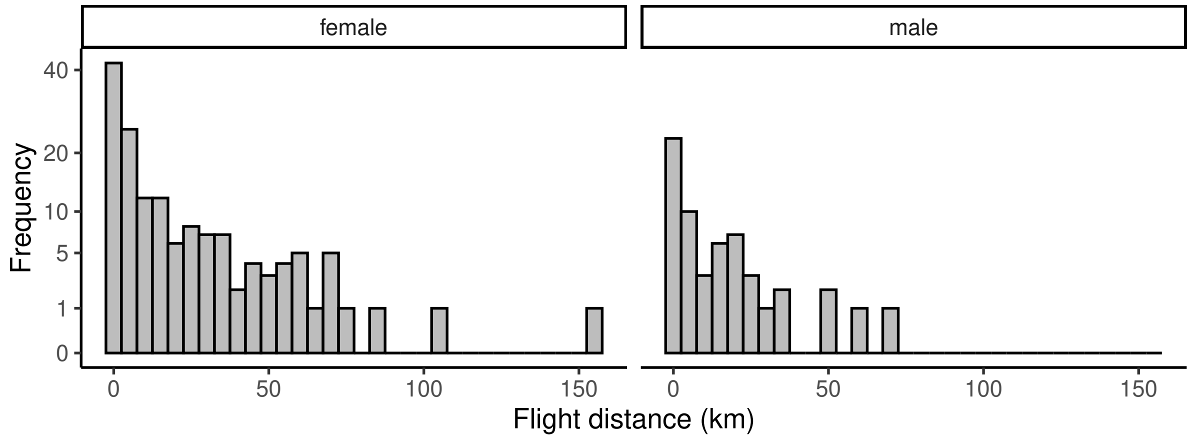


**Figure S3** – Distribution of distances flown by *Cydalima perspectalis* for each sex over the insect lifespan. Experiment 2017. No distinction between modalities and generations was done as their distributions were showing similar shapes. Frequency is displayed with square root scale to improve visibility.

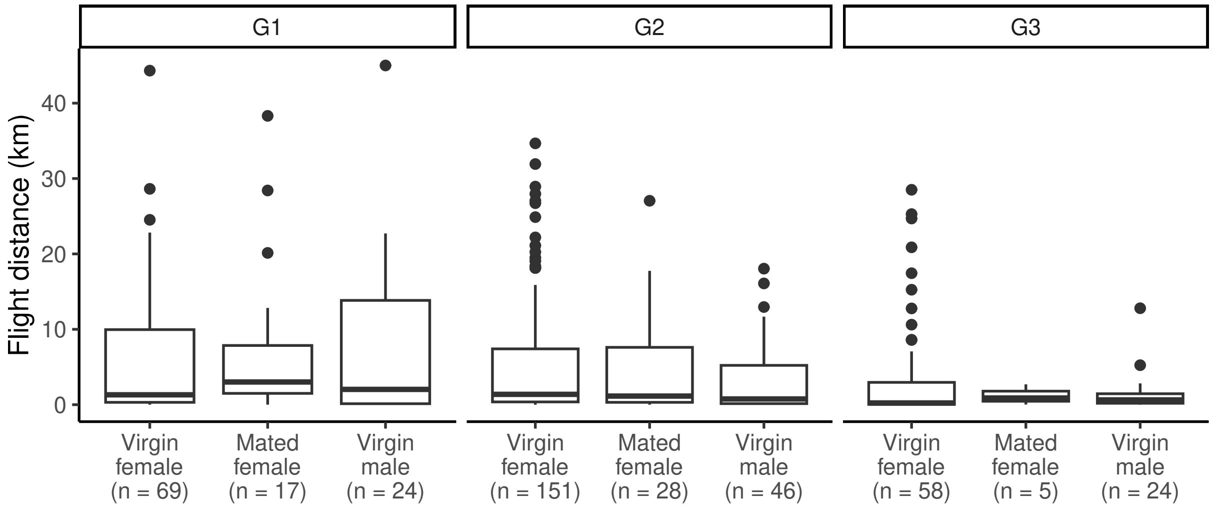


**Figure S4** – Total distances covered by *Cydalima perspectalis* adults during a test under varying mating status. Experiment 2017, A-Normal. Data outside the five first flight tests were discarded to limit age effect. Number of adults in each group is indicated under the group names. In each generation, no significant difference between groups was observed.

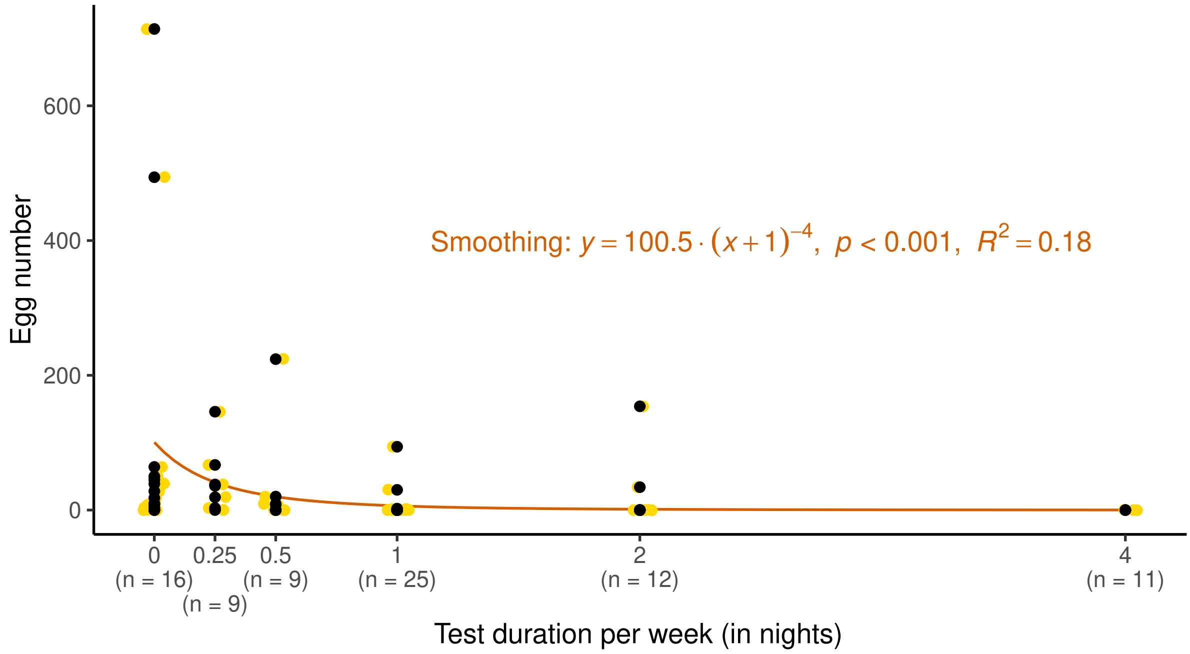


**Figure S5** – Fecundity of *Cydalima perspectalis* females under varying flight test intensity. Experiment 2018. Spearman correlation: $p < 0.001$, $\rho = -0.51$. Jittered gold points partially reveal the position of the overlapping points.
